# SARS-CoV-2 Spike S1 glycoprotein is a TLR4 agonist, upregulates ACE2 expression and induces pro-inflammatory M_1_ macrophage polarisation

**DOI:** 10.1101/2021.08.11.455921

**Authors:** Mohamed M. Aboudounya, Mark R. Holt, Richard J. Heads

**Author notes:** Address for correspondence: Dr R.J. Heads.

## Abstract

**Background and aims:** TLR4 is an important innate immune receptor that recognizes bacterial LPS, viral proteins and other pathogen associated molecular patterns (PAMPs). It is expressed on tissue-resident and immune cells. We previously proposed a model whereby SARS-CoV-2 activation of TLR4 via its spike glycoprotein S1 domain increases ACE2 expression, viral loads and hyperinflammation with COVID-19 disease [1]. Here we test this hypothesis *in vitro* and demonstrate that the SARS-CoV-2 spike S1 domain is a TLR4 agonist in rat and human cells and induces a pro-inflammatory M1 macrophage phenotype in human THP-1 monocyte-derived macrophages.

**Methods:** Adult rat cardiac tissue resident macrophage-derived fibrocytes (rcTMFs) were treated with either bacterial LPS or recombinant SARS-CoV-2 spike S1 glycoprotein. The expression of ACE2 and other inflammatory and fibrosis markers were assessed by immunoblotting. S1/TLR4 co-localisation/binding was assessed by immunocytochemistry and proximity ligation assays on rcTMFs and human HEK-293 HA-TLR4-expressing cells. THP-1 monocytes were differentiated into M1 or M2 macrophages with LPS/IFNγ, S1/IFNγ or IL-4 and RNA was extracted for RT-qPCR of M1/M2 markers and ACE2.

**Results:** TLR4 activation by spike S1 or LPS resulted in the upregulation of ACE2 in rcTMFs as shown by immunoblotting. Likewise, spike S1 caused TLR4-mediated induction of the inflammatory/wound healing marker COX-2 and concomitant downregulation of the fibrosis markers CTGF and Col3a1, similar to LPS. The specific TLR4 TIR domain signalling inhibitor CLI-095 (Resatorvid®), blocked the effects of spike S1 and LPS, confirming that spike S1 is a TLR4 agonist and viral PAMP (VAMP). ACE2 expression was also inhibited by the dynamin inhibitor Dynasore®, suggesting ACE2 expression is mediated by the alternative endosomal/β-interferon pathway. Confocal immunofluorescence microscopy confirmed 1:1 stoichiometric spike S1 co-localisation with TLR4 in rat and human cells. Furthermore, proximity ligation assays confirmed spike S1 and TLR4 binding in human and rat cells. Spike S1/IFN-γ treatment of THP-1-derived macrophages induced pro-inflammatory M_1_ polarisation as shown by an increase in IL-1β and IL-6 mRNA.

**Conclusions:** These results confirm that TLR4 is activated by the SARS-CoV-2 spike protein S1 domain and therefore TLR4 may be a receptor/accessory factor for the virus. By binding to and activating TLR4, spike S1 caused upregulation of ACE2, which may facilitate viral entry into cells. In addition, pro-inflammatory M_1_ macrophage polarisation via TLR4 activation, links TLR4 activation by spike S1 to inflammation. The clinical trial testing of CLI-095 (Resatorvid®) and other TLR4 antagonists in severe COVID-19, to reduce both viral entry into cells and hyperinflammation, is warranted. Our findings likely represent an important development in COVID-19 pathophysiology and treatment, particularly regarding cardiac complications and the role of macrophages.

## INTRODUCTION

The SARS-CoV-2 virus has caused a global COVID-19 pandemic, infecting millions of people worldwide and resulting in millions of deaths. The virus has a spike glycoprotein on its outer envelope, which it uses to enter human cells through interaction with the proposed entry receptor, ACE2 [2, 3]. However, since the pulmonary expression of ACE2 is low (approx. 1-2% of cells in lungs) [4-6], when COVID-19 is primarily a respiratory illness, besides the hyperinflammatory state that occurs in COVID-19 often referred to as the “cytokine storm” [7], has raised questions regarding the pathophysiology of SARS-CoV-2 [1]. We recently proposed a model for SARS-CoV-2 infectivity and disease in which SARS-CoV-2 binds to and activates TLR4 via its spike glycoprotein to increase ACE2 expression, thus facilitating viral entry [1]. Furthermore, the hyperinflammation and myocarditis seen in COVID-19 may be due to over-activation of TLR4 by spike glycoprotein in viral sepsis, leading to an excessive release of pro-inflammatory cytokines downstream of TLR4 [1]. Therefore, in this study, we sought to confirm our original hypothesis in rat and human cells.

Toll-like receptor 4 (TLR4) is a critical pattern recognition receptor (PRR) that not only belongs to the innate immune system but is also expressed on tissue-resident cells as part of intrinsic host defence against invading pathogens [8-10]. Its function is to recognize certain bacterial-, viral-, fungal- and parasitic-derived components, so-called pathogen-associated-molecular-patterns (PAMPs), and direct both the cellular and the systemic immune response [1, 11]. Activation of TLR4 results in the secretion of pro-inflammatory cytokines and chemokines that cause inflammation and alert other cells to infection, in coordination with the priming of adaptive immunity [12, 13]. Activation of TLR4 by viral PAMPs (or VAMPs) results in the secretion of type 1 interferons, such as β-interferon, whose function is to alert neighbouring cells to viral infection, inhibit viral replication and confer an anti-viral state [1, 14, 15]. TLR4 is a very important and extraordinary receptor since it recognizes many different components from different foreign infectious organisms as a first line of defence, in addition to damage-associated molecular patterns (DAMPs) released from host tissue (see [1, 11] and references within). It is a unique receptor that is present on both the cell surface and in endosomes (ie following receptor endocytosis), and is the only TLR having canonical and alternative downstream signalling pathways [16, 17]: (i) a canonical MyD88-dependent pathway which results in the secretion of pro-inflammatory cytokines and chemokines; (ii) an alternative TRIF/TRAM-dependent endosomal pathway which leads to the secretion of type I interferons and some anti-inflammatory cytokines (see [1] and references within). Weaker or delayed (late phase) pro-inflammatory signalling also occurs via the alternative pathway, and there is cross talk between both pathways [18, 19]. However, gram-negative bacterial lipopolysaccharide (LPS), its classical or canonical agonist [20], results in the internalisation of the LPS-TLR4 complex into endosomes within 30 min after the first wave of NFκB activation, followed by signalling mainly in an anti-inflammatory and β-interferon context [13, 21, 22]. The internalization of TLR4 is both dynamin and clathrin-dependent [23].

The SARS-CoV-2 transmembrane spike glycoprotein consists of 2 subunits, S1 and S2 [24, 25]. The outer/external protruding S1 subunit is the most important and was used in this study because it contains the putative binding site to TLR4 based on *in silico* molecular simulations [26], and the Receptor Binding Domain (RBD) which binds to ACE2 [24, 25]. One mechanism called ‘entry by direct fusion’, proposes that after S1 initially binds to ACE2 via the RBD, the S1 subunit is cleaved by the proteases furin and/or TMPRSS2 at 2 cleavage sites (S1/S2 and S2’) [2, 27-31], shed from the virus surface and released into the extracellular space, while the S2 domain then becomes exposed and mediates fusion of the viral and host cell membranes to allow the completion of the viral entry process [24, 27]. Therefore, we hypothesise that S1, either during initial infection, on assembled and released virions or shed from cells during infection, would then activate TLR4 in neighbouring cells.

However, although S1 cleavage and shedding at the cell membrane has been established for SARS-CoV and SARS-CoV-2, for the latter the furin-dependent S1-S2 cleavage may also occur in the trans-Golgi complex during post-translational processing of newly synthesised spike protein post-infection [32-36] (and see [37]). This is because unlike SARS-CoV, SARS-CoV-2 contains an additional unique multi-basic PRRAR motif, which is a furin cleavage site at the junction between S1 and S2 [24, 28]. This distinctive inserted sequence is not present in SARS-CoV and other close viral relatives and would allow it to be cleaved by furin endoproteases, which are ubiquitously expressed in human cells [24, 38-40]. SARS-CoV, MERS-CoV and SARS-CoV-2 utilise both cell surface and endosomal entry pathways and can be primed at the S2’ site by TMPRSS2/4 at the cell surface or by cathepsins B/L in the endosome. Interestingly, a number of entry facilitators or co-factors have been identified including heparan sulphate proteoglycans (HSPGs), phophatidyl serine (PS) receptors, neuropilin-1 (NRP-1), CD147 and C-type lectins (see [37]). Interestingly, the PS receptor T-cell immunoglobulin and mucin domain type-1 (TIM-1) has been shown to enhance the entry of ebola virus via the endosomal route [41, 42]. Therefore, it is intriguing to speculate that TLR4 may be a novel co-factor for SARS-CoV-2 entry via the endosomal pathway.

The present research also sheds light on the implications and significance of the process of spike S1 domain being externalised from cells, either as packaged into new virions or shed as the cleaved S1 domain (see [32] and refs within). We hypothesise that spike S1 would bind and activate TLR4 to increase ACE2 expression and contribute to the release of pro-inflammatory cytokines.

We sought to confirm our hypothesis in adult rat cardiac tissue resident macrophage-derived cells (cTMFs) which we routinely use in our research and which co-express monocyte/macrophage and myofibroblast markers and adopt a myofibroblast (fibrocyte) phenotype in culture and play an important role in myocardial inflammation and wound healing [9]. These cells also have a low basal ACE2 expression; unlike cardiomyocytes where at least 7.5% may be ACE2 positive [4]. More importantly, cTMFs express TLR4 and respond to LPS and DAMPs such as the extra domain A of fibronectin (FN_EDA_) [9]. In addition, we used human cells expressing TLR4 (hTLR4-HA HEK 293 cells) and human monocyte-derived macrophages which also express TLR4.

Here we show that SARS-CoV-2 spike S1 subunit binds and activates TLR4 as well as upregulates ACE2 expression in rat cTMFs as shown by immunoblotting, RT-qPCR, confocal immunofluorescence microscopy and proximity ligation assay. Furthermore, spike S1 caused similar downstream effects as bacterial LPS on the expression of certain inflammatory and fibrotic markers, including cyclo-oxygenase-2 (COX-2), connective tissue growth factor (CTGF) and the collagen 3a1 isoform (COL3A1). The effects of spike S1 were inhibited by the selective TLR4 signalling inhibitor CLI-095 (Resatorvid® or TAK-242), confirming that spike S1 is a TLR4 agonist and a viral PAMP for TLR4. Proximity ligation assays confirm spike S1 binding to TLR4 in both rat cTMFs and human TLR4 expressing cells. Treatment of human THP-1 differentiated macrophages with spike S1 and Interferon-γ (IFNγ) resulted in the induction of a pro-inflammatory M1 macrophage phenotype, comparable to that induced by bacterial LPS plus IFNγ. Thus, TLR4 activation likely contributes to the hyperinflammatory state observed with SARS-CoV-2 viral sepsis. Overall, we have confirmed our hypothesis and our findings also highlight the role of macrophages in COVID-19 and provide possible explanations as to why the virus is so virulent and can cause severe multi-organ disease.

## MATERIALS AND METHODS

### Materials

General-purpose chemicals and reagents were from Thermo Fisher Scientific UK, and Sigma-Aldrich UK, unless specified otherwise. General cell culture media and reagents were from Sigma-Aldrich, UK and Gibco, UK. Fetal Bovine Serum (FBS: sterile filtered Australian Origin Bovine Serum) was obtained from Pan Biotech (P40-39500); LPS (Lipopolysaccharide) was *Salmonella enterica* serotype typhimurium (L2262) from Sigma-Aldrich UK; CLI-095 (Resatorvid®; TAK242) was from Sigma-Aldrich, UK; Dynasore® (ab120192) was from Abcam, UK; Recombinant SARS-CoV-2 Spike S1 active glycoprotein, aa16-685 (ab273068), was from Abcam, UK; Recombinant SARS-CoV-2 (N501Y) Spike RBD, aa391-451 protein (40592-V08H82), was from Sino-Biological, USA; Phorbol 12-myristate 13-acetate (PMA) (1201/1) was from Bio-Techne, UK; Cytokines: interferon gamma (IFNγ) (SRP3058) and human Interleukin-4 (IL-4) (I4269) were from Sigma-Aldrich, UK; Primary antibodies were as follows: anti-CTGF goat polyclonal antibody (sc-14939), and anti-COL3A1 goat polyclonal antibody (sc-8781) were from Santa Cruz Biotechnology, USA; anti-COX-2 (PA5-32366) and anti-TLR4 (MA5-16216) were from Invitrogen, UK; anti-ACE2 (ab15348) and anti-spike S1 (ab273074) were from Abcam, UK; anti-GAPDH (14C10), and anti-beta-actin (4970S) were from Cell Signalling Technology, NL. Secondary antibodies were as follows: polyclonal rabbit anti-goat Immunoglobulins/HRP (P0160) and polyclonal swine anti-rabbit Immunoglobulins/HRP (P0217) were from Dako, DK; Cy™3-conjugated AffiniPure goat anti-rabbit IgG (111-165-144) and Cy™3-conjugated AffiniPure goat anti-mouse IgG (115-165-146) were from Jackson Immuno Research Laboratories Inc., USA; Donkey anti-rabbit Alexa Fluor® 647 conjugated IgG (A31573) was from Invitrogen, UK; goat anti-mouse IgG Alexa Fluor® 488 (ab150117) and Goat ant-rabbit Alexa Fluor® 647 conjugated (ab150091) were from Abcam, UK; wheat germ agglutinin (WGA) conjugated to Alexa Fluor® 488 (W11261) was from Thermo Fisher Scientific, UK. Bovine serum albumin (BSA) fraction V was from Roche diagnostics, GmbH; borosilicate glass coverslips were from VWR, UK; Vector-Shield hard-set mounting medium with DAPI was from Vector Labs, USA; Page Ruler™ Prestained protein ladder (26616) was from Thermo Fisher Scientific, UK; pre-cast 10% mini-Protean® TGX SDS-PAGE mini-gels were from Biorad, UK; polyvinylidene difluoride (PVDF) membranes (Hybond-P) were from Amersham, UK; Enhanced Chemiluminescence Western Blotting Substrate (ECL) and the CL-Xposure film were from Thermo Fisher Scientific, UK; Monarch® total RNA miniprep kit was from New England Biolabs, UK; PCR primers were from Insight Biotechnology, UK, Integrated DNA Technologies, BE and Invitrogen, UK.

### Isolation of Rat cTMFs

cTMFs were isolated from adult rat ventricles by collagenase digestion on a Langendorff perfusion apparatus. The non-myocyte cell fraction containing tissue resident macrophages was obtained as the supernatant following the settling of myocytes through BSA and centrifuged at 1000 rpm for 5 min to pellet the cells, followed by resuspension in Full Growth Medium (FGM: Dulbecco’s Modified Eagle’s Medium-high glucose (DMEM); 1% penicillin/streptomycin; 10% foetal calf serum). Cells were washed x2 by pelleting at 1000 rpm for 5 min followed by resuspension in FGM. Cells were finally diluted in FGM at the appropriate volume then plated onto polystyrene 6-well plates, grown in FGM followed by a media change 2-3 days later, then after reaching confluence, media was changed to serum free (SF) media 24 hours prior to treatment with agonists. cTMFs were characterized as co-expressing monocyte/macrophage, mesenchymal stem cell and myofibroblast markers including CD90 (Thy1); F4/80; CD14; Mrc1; Col1a1; Col3a1; alpha-smooth muscle actin (αSMA); vimentin; periostin etc elsewhere [9] and manuscript in preparation.

### Pharmacological treatment

Cells were treated with TLR4 agonists, either LPS (2 μg/ml) or Spike S1 (100ng/ml) in 2 ml medium per well on 6-well plates. Inhibitors were added 1 hour prior to agonist addition. CLI-095 was used at 2.8 μM final concentration (from a 1mg/ml stock solution in DMSO. Final DMSO concentration was 0.1%). Dynasore® (dynamin inhibitor) was used at 80 µM final concentration. The cells were harvested approximately 22 hours later. For cytokine treatment on THP-1 cells, LPS (50 ng/ml) plus IFNγ (50ng/ml); Spike S1 (100ng/ml) plus IFNγ (50 ng/ml); IFNγ alone (50 ng/ml), IL-4 (20 ng/ml), as outlined in the section below (THP-1 cells and differentiation).

### Sample preparation

Cells were lysed in 200 µl of 2x SDS-PAGE sample buffer (20% glycerol; 6% SDS in 0.12 M Tris, pH 6.8) per well, and scraped into microcentrifuge tubes. 2-mercapto-ethanol was added to the samples (10% vol/vol). Samples were heated at 100°C for approx. 7 minutes, followed by cooling at 4°C. 4μl of Bromophenol blue (8% in ethanol) was then added to each sample. Samples were then stored at -20°C prior to analysis on 10% SDS-PAGE gels.

### SDS-PAGE and Immunoblotting

Samples were run on pre-cast mini-Protean® TGX 10% SDS-PAGE gels (Bio-Rad, UK) using a mini-protean II apparatus (Bio-Rad, UK) at a constant 85V. Proteins were then electrophoretically transferred onto PVDF membranes overnight at 30mA, via wet transfer using Mini Trans-Blot Cell (Bio-Rad, UK). Following transfer, membranes were blocked in a solution containing PBS; 0.05% (v/v) Tween 20; 2% (w/v) milk powder for 1 hour at room temperature. Membranes were then incubated in primary antibody at a 1:1000 dilution in PBS; 0.05% Tween 20; 0.1% milk solution (PBSTM) for approx. 3 hours at room temperature with constant gentle shaking. The membranes were then washed for 3x for 5 min each in PBSTM. The corresponding horseradish peroxidase (HRP)-conjugated secondary antibodies (Dako, Denmark), diluted at 1:2500 in PBSTM were added for approx. 1 hour at room temperature. Membranes were then washed in PBSTM 3x for 5 min each. They were then developed by exposing the membranes to enhanced chemiluminescence (ECL) detection reagent and exposing them to CL-Xposure film in a cassette in the dark for between 30 s and 2 min. Films were developed using an RGII X-Ray Film Processor (Fuji, Japan).

### Immunocytochemistry & Confocal fluorescence microscopy

Briefly, cells were grown on sterilized borosilicate glass coverslips in 6 well plates to approx. 80% confluence for approximately 1 week. Media was changed to serum free for 24 hours, then LPS (2µg/ml) or Spike (100ng/ml) were added as appropriate to the experiment, or left untreated as controls, and the cells were incubated for a further 22 hours. Cells on coverslips were washed with 1 ml of cold PBS and fixed with 1 ml of 4% paraformaldehyde for 10 minutes at room temperature. The coverslips were washed twice with 1 ml of cold PBS for five minutes each. Cells were then permeabilized with 1ml of 0.2% Triton X-100 in PBS for 20 min at room temperature and washed twice with 1ml of cold PBS for five minutes each. Following this, they were incubated for two hours at room temperature in 1 ml of blocking buffer (0.1% Triton X-100 and 2% Bovine Serum Albumin (BSA) in PBS). This was followed by adding 800 µl (final volume) of diluted primary antibody in 2% BSA in PBS, to the coverslips as appropriate to the experiment and incubated overnight at 4°C. The antibody concentration varied and depended on the antibody used, but it was usually at 1:200 or similar for the primary antibodies (i.e. TLR4) and 1:400 for ACE2. Coverslips were then washed for 3x 5 minutes each in 1 ml of cold PBS. 800 µl of the relevant fluorescent secondary antibodies were added, diluted in 2% BSA in PBS and incubated in the dark for 2 hours at room temperature. This was followed by 3x washes for 5 minutes each in 1 ml of PBS at room temperature and allowing it to air dry. The coverslips were then mounted on glass slides, using mounting medium with DAPI. Edges of coverslips were sealed with nail varnish and left to set. Slides are stored at 4°C in the dark. The slides were analysed using a Leica TCS SP5X confocal microscope. Confocal images were analysed at x63 or x100 magnification.

### Ratiometric analysis of S1 spike and TLR4 signal overlap

16-bit confocal images were imported into Wolfram Mathematica 12.3 (Champaign II), denoised and background subtracted using a Difference of Gaussians with kernel radii of 2 and 127 pixels respectively. The arctangent of the red channel intensity with respect to the green channel intensity was then calculated per pixel and normalised to a scale of 0 - 1. This is a linear scale that gives the degree of overlap between two fluorophores. Values closer to 0 are due to a higher presence of red fluorophore per pixel, whilst values closer to 1 have a higher presence of green fluorophore. Values of 0.5 represent an equal presence of each fluorophore. A histogram of these ratios was then plotted using the Norm of the two intensities as a weight. The same weighting was also used to calculate the weighted mean of the distribution.

### Proximity ligation assay

Proximity ligation assay (PLA) was performed using the Duolink® In Situ Red Mouse/Rabbit kit (DUO92008, Sigma-Aldrich, UK) according to the manufacturer’s instructions. Primary antibodies used were mouse anti-human TLR4 and rabbit anti-spike S1 with the appropriate secondary antibodies (anti-rabbit-minus and anti-mouse-plus PLA probes) from the kit and red detection reagents.

### 293-hTLR4-HA cells

293-hTLR4-HA cells (InvivoGen, USA) are HEK 293 cells stably transfected with the hTLR4a gene fused at the 3’ end to an influenza haemagglutinin (HA) tag. 293 cells express very low levels of endogenous TLR4. Cells were maintained in FGM in the presence of normocin™ (100ug/ml) and the selective antibiotic blasticidin S (10 ug/ml). 293 Null cells were used as a negative control.

### THP-1 cells and differentiation

The human monocytic cell line THP-1 was routinely maintained in RPMI 1640 (1x) growth medium containing 10% of heat-inactivated FBS, 4.5 mg/ml D-glucose, 2mM L-glutamine, 10mM HEPES, 1mM pyruvate, 0.05 mM 2-mercaptoethanol, and 1% Penicillin/streptomycin. Cells were differentiated into M_ø_ macrophages by treatment with phorbol myristyl acetate (PMA, 80nM) for 48 hrs, followed by recovery for 24 hours. M_ø_ macrophages were then further differentiated into M_1_ macrophages by treatment with LPS plus IFNγ (50ng/ml +50ng/ml), or Spike S1 plus IFNγ (100 ng/ml + 50 ng/ml) for 24 hrs or into M_2_ macrophages by treatment with IL-4 (20 ng/ml) for 24 hrs [43].

### RNA extraction and RT-qPCR

Samples were isolated in RNA extraction buffer and RNA isolated using the Monarch® Total RNA extraction Miniprep kit (T2010S; New England Biolabs) according to the manufacturer’s instructions. RNA concentration and yield were measured using a ThermoScientific™ Nanodrop 2000 spectrophotometer. Then, 500ng RNA was reverse transcribed using the High Capacity RNA-to-cDNA kit (4387406, Applied Biosystems, UK), and RT-qPCR was carried out on the cDNA samples using the following primers:

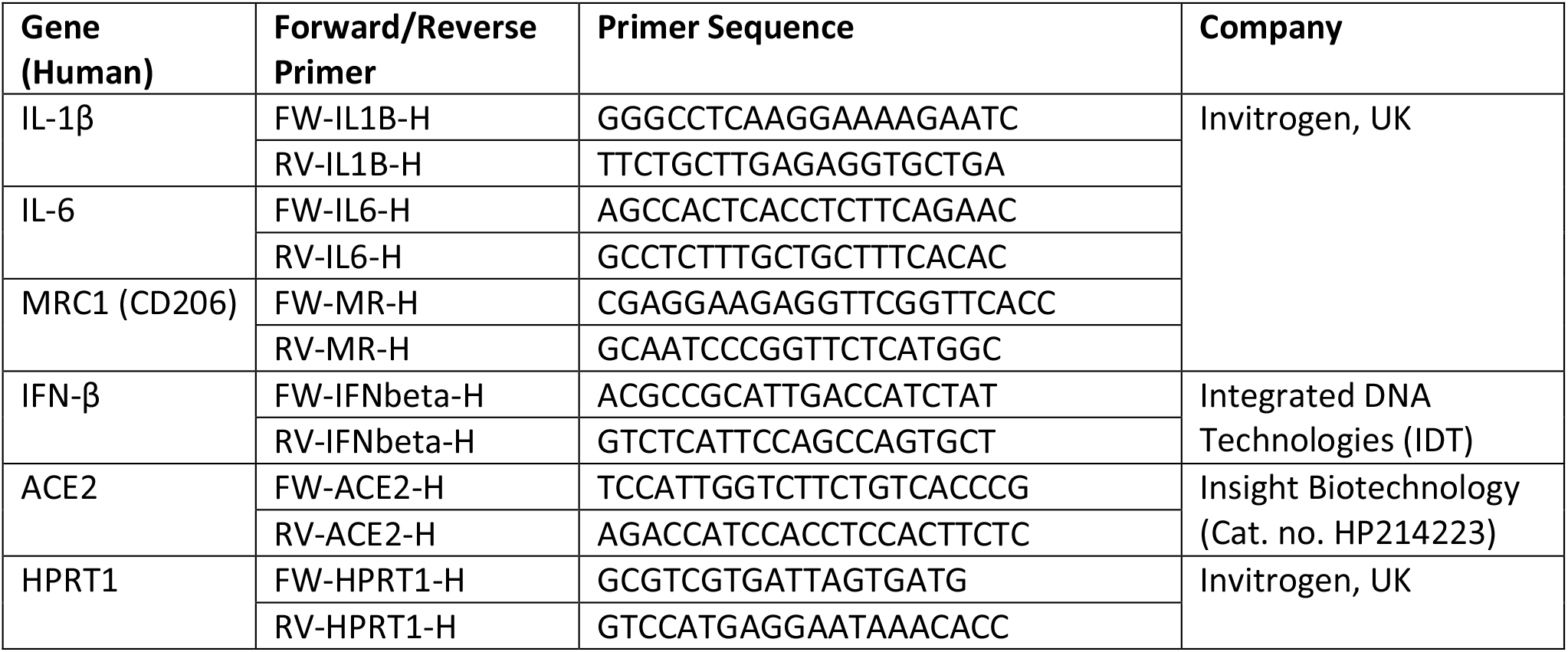

RT-qPCR was carried out in a 20uL reaction mixture per well using the PowerUp™ SYBR™ green Master Mix (A25742; Applied Biosystems, UK) in MicroAmp Fast Optical 96-well plates (0.1 ml) (4346907; Applied Biosystems, UK) on an ABI Prism 7500 Fast qPCR machine. Cycle conditions were 95°C for melting and 60°C for annealing for 40 cycles.

### Statistical analysis

Immunoblots were scanned using a desktop scanner and band densities analysed using Image Studio Lite™ (LiCor®). Protein of Interest (POI) integrated band densities were normalised to the corresponding GAPDH band and expressed as a ratio. Statistical analysis was performed and graphs were produced using GraphPad Prism 8, either using a Kruskal Wallis test, Wilcoxon Signed Rank Test, or Repeated Measures One-way ANOVA with G-G correction followed by Tukey’s multiple comparisons post hoc test as appropriate to the experiment and as indicated in the figure legends. Data is expressed as box plots with median and full range. For RT-qPCR data was expressed as mean ± S.E.M as dot plots.

## RESULTS

### SARS-CoV-2 spike S1 glycoprotein causes upregulation of ACE2

Under baseline conditions ACE2 expression in cTMFs was low. Figure 2 shows that treatment of the cells with spike S1 glycoprotein led to the prominent induction of ACE2 expression to 2-3 fold over control levels, at approx. 22 hours following treatment. In addition, LPS induced ACE2 protein expression approx. 4-fold at the same time point. Hence, it is apparent that TLR4 activation by either LPS or spike S1 causes the upregulation of ACE2. The concentration of spike S1 used (100 ng/ml or 0.8nM) was below concentration range that has been shown to bind to and activate ACE2 (1.2-14.7nM) [24, 25], suggesting stronger binding to TLR4 and as predicted by molecular docking/binding enthalpy predictions [26].

**Figure 1.**
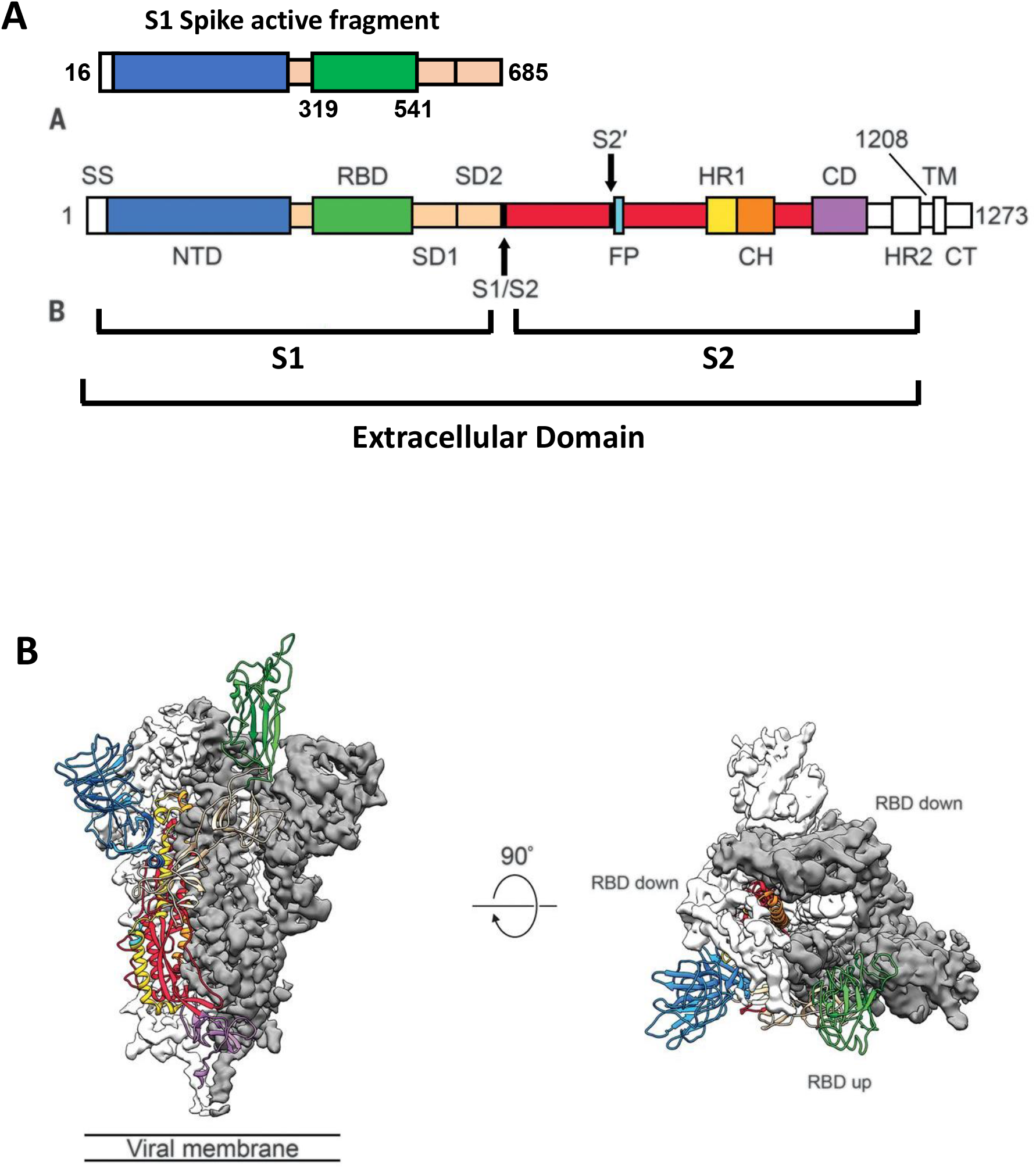
Structure of SARS-CoV-2 spike protein. **Panel A:** Schematic representation of the SARS-CoV-2 spike protein primary structure. The S1 spike protein active fragment (aa 16-685) as used in the experiments is shown at the top with the receptor binding domain (RBD: aa 319-541) used as a control, indicated. The N-terminal domain (NTD) is shown in blue and RBD in green. Below is shown the full-length spike protein with the S1 and S2 subunits. NTD (blue); RBD (green); S1/S2 boundary [protease cleavage site (arrow); S2’ protease cleavage site (arrow); structural domains/linker region (SD1 and SD2: beige); fusion peptide (FP: sky blue); heptad repeat 1 (HR1: yellow); Central helix (CH: orange); Connector domain (CD: purple); the N-terminal Signal sequence (SS), HR2, transmembrane segment (TM) and C-terminal domain cytoplasmic tail (CT) are show in white. **Panel B:** The prefusion structure of the SARS-CoV-2 spike protein complex. Side view (left) and top view (right) with a single RBD in the up (open) conformation. In this depiction the two RBD down protomers are shown as cryo-EM density maps in white or grey and the RBD up protomer is shown as a ribbon diagram (green). Adapted from Wrapp et al [25] (with permission).

**Figure 2.**
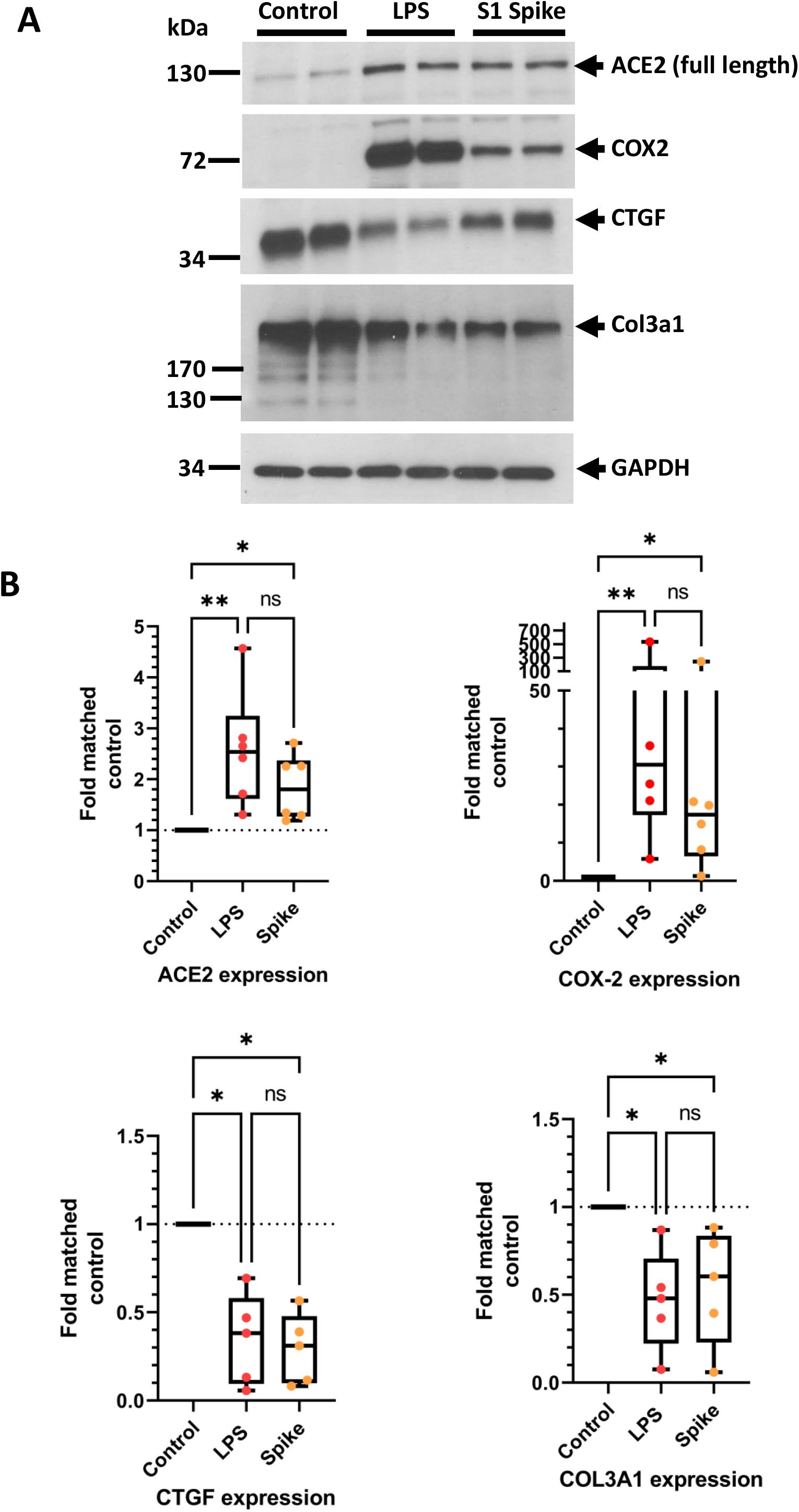
S1 Spike induced activation of TLR4. **Panel A:** Cultured primary rat cardiac tissue macrophage-derived fibrocytes (cTMFs) were treated with gram -ve bacterial lipopolysaccharide (LPS) (2 μg/ml) or recombinant S1 spike protein (aa 16-685) (100 ng/ml) and harvested 22 hours later for immunoblot analysis. Blots were probed with antibodies against the SARS-CoV-2 entry receptor ACE2 and signature markers of the typical response of these cells to TLR4 activation [9]: cyclooxygenase-2 (COX-2), connective tissue growth factor (CTGF/CCN2) and collagen isoform 3a1 (Col3a1). Blots were also probed for GAPDH as a loading control. The position of molecular weight markers in kDa are shown on the left for comparison. **Panel B:** Quantitative analysis of ACE2, COX-2, CTGF and Col3a1 protein expression. Data for each protein was normalised to the corresponding GAPDH band and expressed as fold matched untreated control set to 1. Data for each protein of interest (POI) are expressed as box plots with median, inter-quartile range and full range. Statistical significance is shown as *p≤0.05 or **P≤0.01 following a Kruskal-Wallis test.

### SARS-CoV-2 S1 glycoprotein is similar to LPS with respect to regulation of downstream gene expression

We have previously shown that TLR4 activation by LPS and the DAMP fibronectin extra domain A (FN_EDA_) in rat cTMFs causes the induction of COX-2 expression and reciprocal downregulation of the fibrosis markers connective tissue growth factor (CTGF) and the collagen 1a1 and 3a1 isoforms [9]. Indeed, spike S1 glycoprotein caused a similar, but weaker, induction of COX-2 expression and downregulation of CTGF and Col3a1 expression in a similar manner to LPS (Figure 2), therefore inducing a signature response to TLR4 activation in these cells comparable to LPS. These findings confirm that spike S1 is a viral PAMP (VAMP) for TLR4.

### SARS-CoV-2 spike S1 glycoprotein is a TLR4 agonist and induces the expression of ACE2 via the alternative endosomal pathway

In order to confirm that spike S1 glycoprotein subunit is an agonist for TLR4, we tested its effects in the presence of the specific, cell-permeable TLR4 signalling inhibitor CLI-095 (also known as Resatorvid® or TAK-242). This drug selectively binds to the TLR4 intracellular TIR domain, thereby inhibiting the interactions between TLR4 and its adaptors [1, 44, 45]. Therefore, CLI-095 inhibits both the canonical MAL/MyD88 and the alternative TRAM/TRIF signalling pathways that are downstream of TLR4. Figure 3, panels A and C (i) and (ii) show that there was a marked increase in ACE2 expression in both LPS-treated and spike S1-treated cells. However, this effect was completely inhibited in the presence of CLI-095.

**Figure 3.**
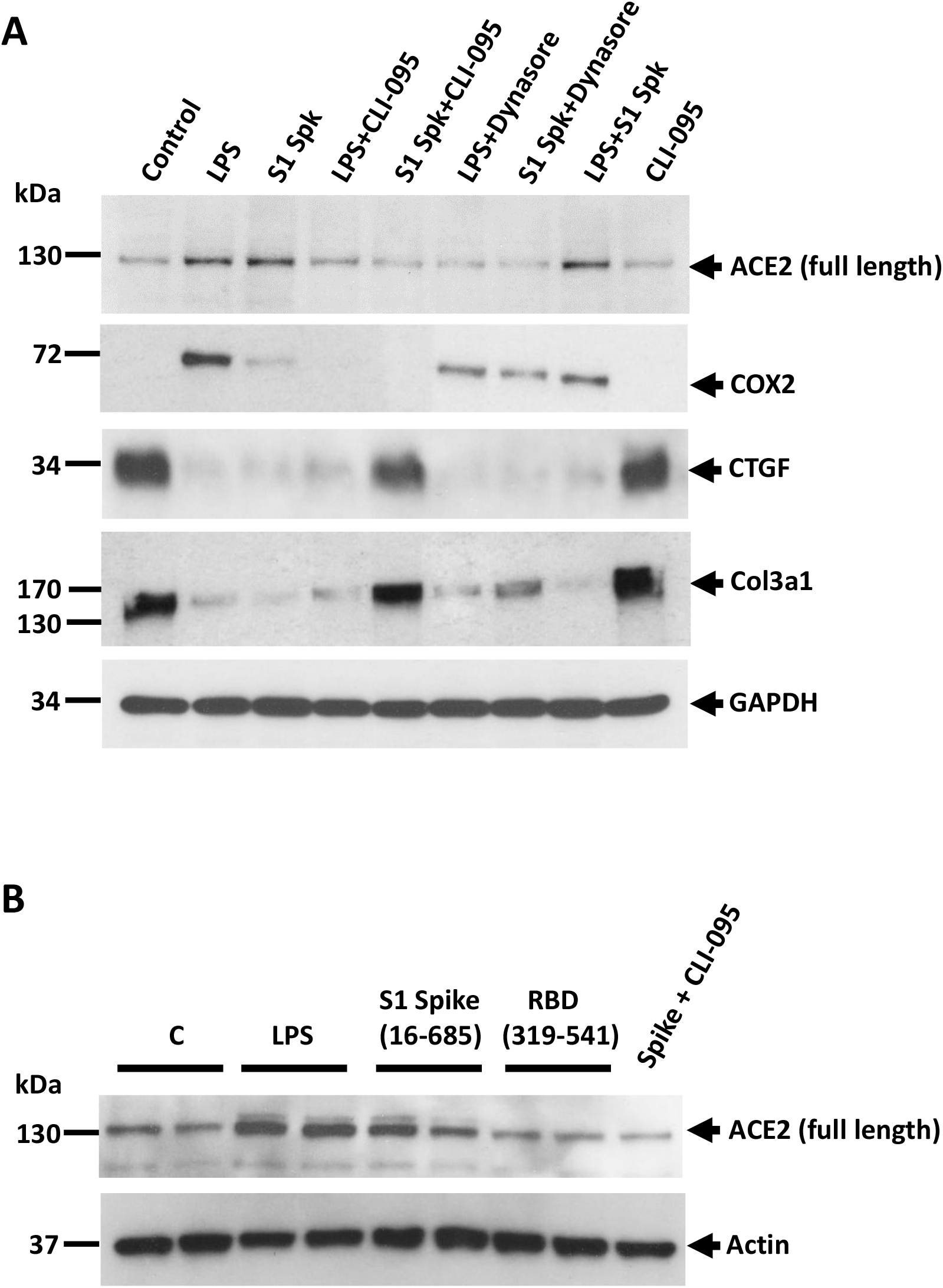
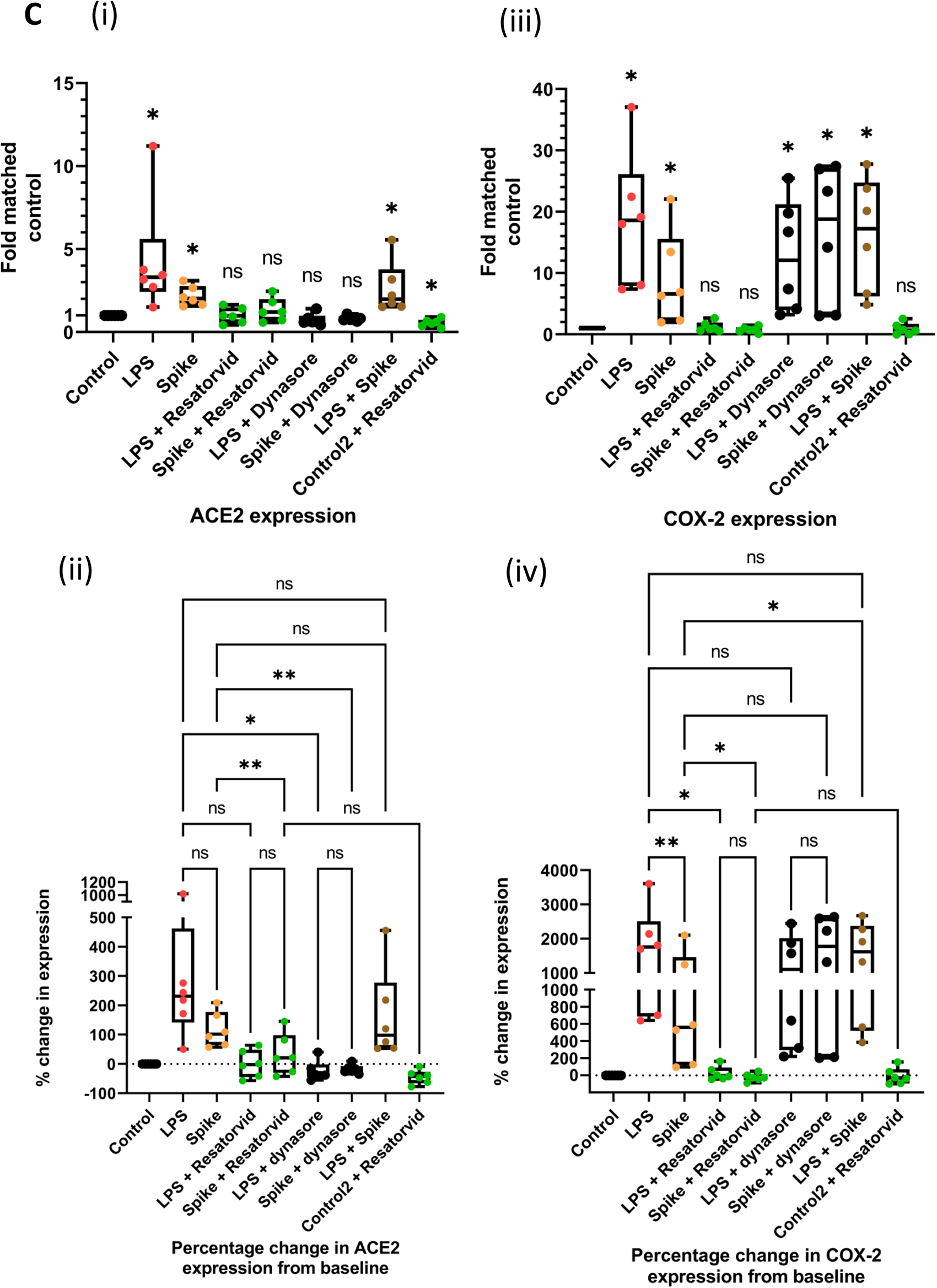
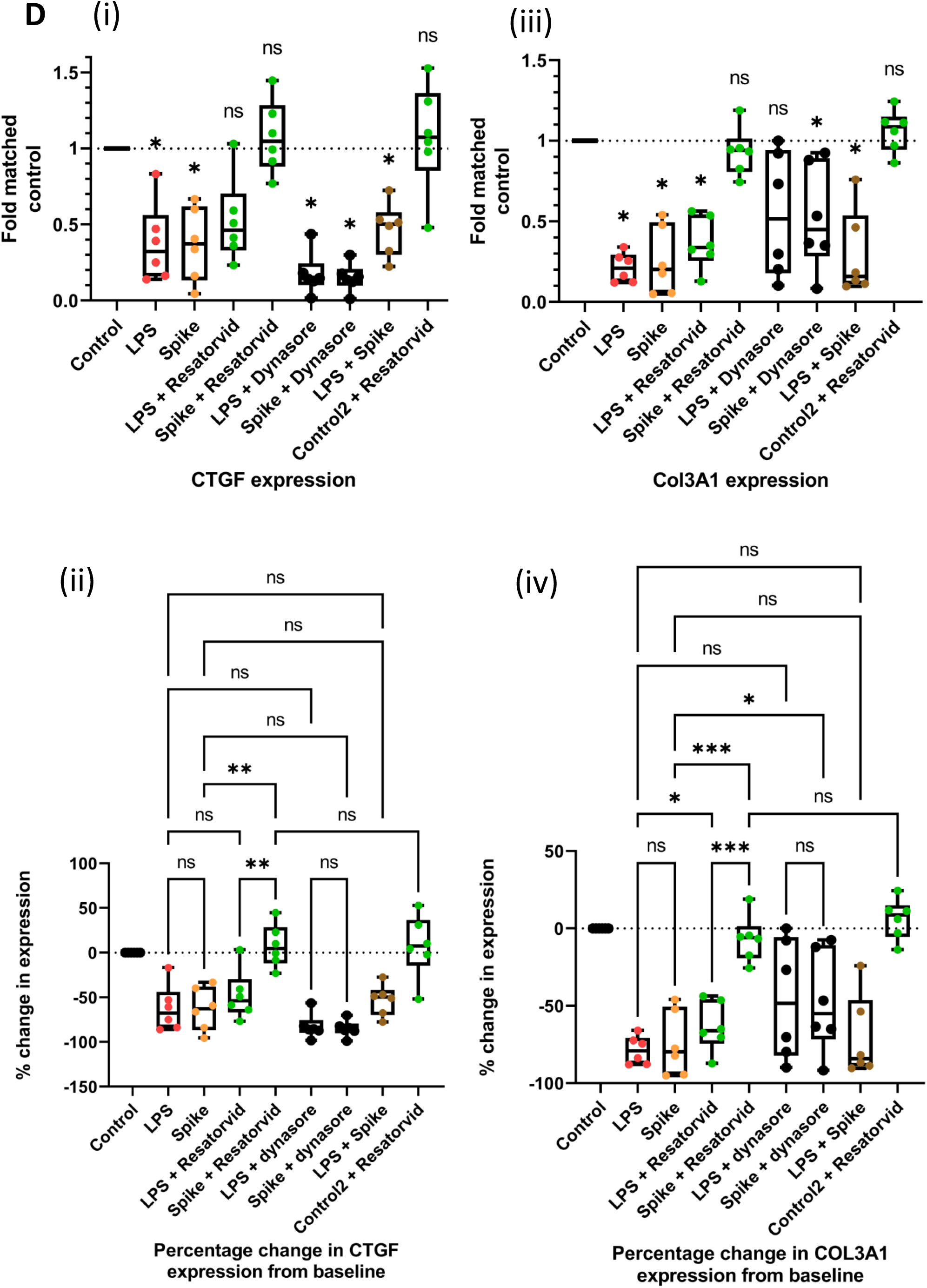
The TLR4 antagonist CLI-095 (Resatorvid®) inhibits S1 spike-induced activation of TLR4. **Panel A:** Cultured primary rat cardiac tissue macrophage-derived fibrocytes (cTMFs) were treated with LPS (2 mg/ml) or recombinant S1 spike protein (100 ng/ml) either alone or following pre-treatment with the TLR4 antagonist CLI-095 (2.8 μM) or the dynamin inhibitor Dynasore™ (80 μM). Cells were harvested 22 hours later for immunoblot analysis. Blots were probed with antibodies against ACE2, COX-2, CTGF and Col3a1. Blots were also probed for GAPDH as a loading control. The position of molecular weight markers in kDa are shown on the left for comparison. **Panel B:** Cultured primary rat cardiac tissue macrophage-derived fibrocytes (cTMFs) were treated with LPS (2 mg/ml), recombinant S1 spike protein aa 16-685 (100 ng/ml) or recombinant S1 spike protein RBD aa 319-541 (100 ng/ml). Cells were harvested 22 hours later for immunoblot analysis and blots were probed with antibody against ACE2. Blots were also probed for β-actin as a loading control. **Panel C:** Quantitative analysis of ACE2 and COX-2 protein expression. Data for each protein was normalised to the corresponding GAPDH band and expressed as fold matched untreated control set to 1. Data for each protein of interest (POI) are expressed as box plots with median, inter-quartile range and full range (n=6). Statistical significance is shown as *p≤0.05 following a Wilcoxon Signed Rank test [sub-panels (i) and (iii)]. For multiple comparisons between groups data was expressed as percent change relative to control and analysed using a Repeated Measures One-Way ANOVA with a Tukey’s post-hoc test (n=6) where statistical significance is shown as *p≤0.05 or **P≤0.01. Data for each protein of interest (POI) are expressed as box plots with median, inter-quartile range and full range [sub-panels (ii) and (iv)]. **Panel D:** Quantitative analysis of CTGF and Col3a1 protein expression. Data for each protein was normalised to the corresponding GAPDH band and expressed as fold matched untreated control set to 1. Data for each protein of interest (POI) are expressed as box plots with median, inter-quartile range and full range (n=6). Statistical significance is shown as *p≤0.05 following a Wilcoxon Signed Rank test [sub-panels (i) and (iii)]. For multiple comparisons between groups data was expressed as percent change relative to control and analysed using a Repeated Measures One-Way ANOVA with a Tukey’s post-hoc test (n=6) where statistical significance is shown as *p≤0.05 or **P≤0.01. Data for each protein of interest (POI) are expressed as box plots with median, inter-quartile range and full range [sub-panels (ii) and (iv)].

Furthermore, pre-treatment with Dynasore®, a specific inhibitor of dynamin i.e. a drug which prevents endosome formation and hence inhibits the TLR4 alternative pathway, also resulted in inhibition of ACE2 induction. Dynasore® inhibits endocytosis in macrophages and therefore specifically inhibits early endosomal TRIF-dependent TLR4 signalling [46, 47]. This suggests that spike S1 upregulates ACE2 via TLR4-dependent signalling and specifically via the alternative endosomal pathway. Treatment of cells with LPS plus spike together also caused an increase in ACE2 expression, but this was not additive, suggesting maximal activation in the presence of LPS alone. Control cells treated with CLI-095 alone had no effect on basal ACE2 levels or cell viability as shown by GAPDH levels.

To confirm that the effects of spike S1 were specific and not due to a non-specific effect of addition of exogenous recombinant protein to the cells, we used the spike ACE2 receptor binding domain (RBD) as a negative control. In silico molecular simulations have suggested that TLR4 binding involves amino acid residues in the N-terminal domain (NTD-shown in blue in figure 1) and the linker region (shown in beige in figure 1) and not the RBD (shown in green in figure 1) [26]. Figure 3, panel B shows that there was no induction of ACE2 by the RBD alone.

### CLI-095 and/or Dynasore™ block other TLR4-mediated effects of SARS-CoV-2 spike S1 glycoprotein

#### Effects on COX-2 expression

CLI-095 also completely inhibited the induction of COX-2 expression by either LPS or spike S1, confirming TLR4 involvement. Dynasore® partially inhibited COX-2 expression in LPS-treated cells (Figure 3, panel A), but this was not borne out statistically [Figure 3, panel C (iii) and (iv)], suggesting that COX-2 expression is not significantly induced via the alternative TRIF/TRAM endosomal pathway. This suggests that either the MyD88 canonical pathway is mainly responsible for COX-2 induction, given that the COX-2 proximal gene promoter has transcription factor binding sites for NFκB [48, 49], or as we have previously shown in these cells, that COX-2 expression downstream of TLR4 activation is mediated via the calcineurin (CN)/NFAT transcription factor pathway [9].

Dynasore® appeared to potentiate spike S1-mediated COX-2 induction, which suggests that spike is also acting via the MyD88 canonical or CN/NFAT pathway. Blocking the endosomal pathway and the internalisation of TLR4 could increase cell surface expression of TLR4 and enhance signalling via the canonical pathway, especially since the alternative pathway is considered to act as a negative feedback regulator on the canonical pathway through receptor internalisation into endosomes for recycling and the production of some anti-inflammatory cytokines (Figure 3, panel C). CLI-095 alone had no effect on basal COX-2 expression.

#### Effects on CTGF and Col3a1 expression

The expression of the fibrosis markers CTGF and Col3a1 are basally high in cTMFs under normal culture conditions due to their predominantly myofibroblast phenotype, and expression is downregulated by TLR4 activation as previously described [9]. CLI-095 inhibited the downregulation conferred by spike S1, restoring both CTGF and Col3a1 expression to baseline levels similar to control cells (Figure 3, panels A and D). However, CLI-095 incompletely inhibited the downregulating effect of LPS with regards to Col3A1 and to some extent with CTGF. This suggests that there may be slight differences between LPS and spike S1 signalling downstream of TLR4 or that the dose of CLI-095 used was insufficient to completely inhibit LPS. Other co-receptors, particularly CD14 are involved in LPS signalling via TLR4 and also can initiate signalling culminating in NFAT-dependent transcription of genes independently of TLR4 [22, 50]. This would be consistent with our previous finding that NFAT mediated the transcriptional effects on gene expression in these cells [9]. Moreover, CD14 in particular plays a role in bringing LPS to TLR4 [51]. This should be investigated further, but the most important finding of relevance here is that spike S1-mediated effects are inhibited by CLI-095, confirming that it is an agonist for TLR4.

Dynasore® enhanced the downregulation of CTGF caused by either LPS or spike S1. However, Dynasore® partially inhibited the downregulation of Col3a1 by LPS and S1, which suggests complex differences between the regulation of CTGF and Col3a1 expression. CTGF and Col3a1 are secreted proteins, and dynamin is also important for their exocytosis [52, 53]. The canonical or another pathway could also be responsible for downregulation. CLI-095 alone had no significant effect on basal CTGF or Col 3a1 levels.

### Spike S1 glycoprotein co-localization with TLR4

To confirm whether spike S1 co-localisation with TLR4 on cells may be indicative of TLR4 binding, we undertook confocal immunofluorescence microscopy of both spike glycoprotein and TLR4. Figure 4, panel B shows that both TLR4 (red) and spike S1 (yellow) displayed a punctate appearance and significant co-localisation as shown by orange on the overlayed images. Images taken at a higher magnification (x100) confirmed a high degree of overlap of the two fluorophores. To confirm overlap we performed ratiometric analysis of the confocal images. In Figure 5 a pseudo-colour spectrum was applied where red depicts a low degree of co-localisation and cyan depicts high degree of co-localisation, with a ratio of 0.5 indicating 1:1 co-localisation (see materials and methods). It can be clearly seen that in control cells in the absence of spike S1 the TLR4 signal is red, whereas in the presence of spike S1 there is a strong shift towards the cyan and 0.5, with an average ratio of 0.48, suggesting strong co-localisation.

**Figure 4.**
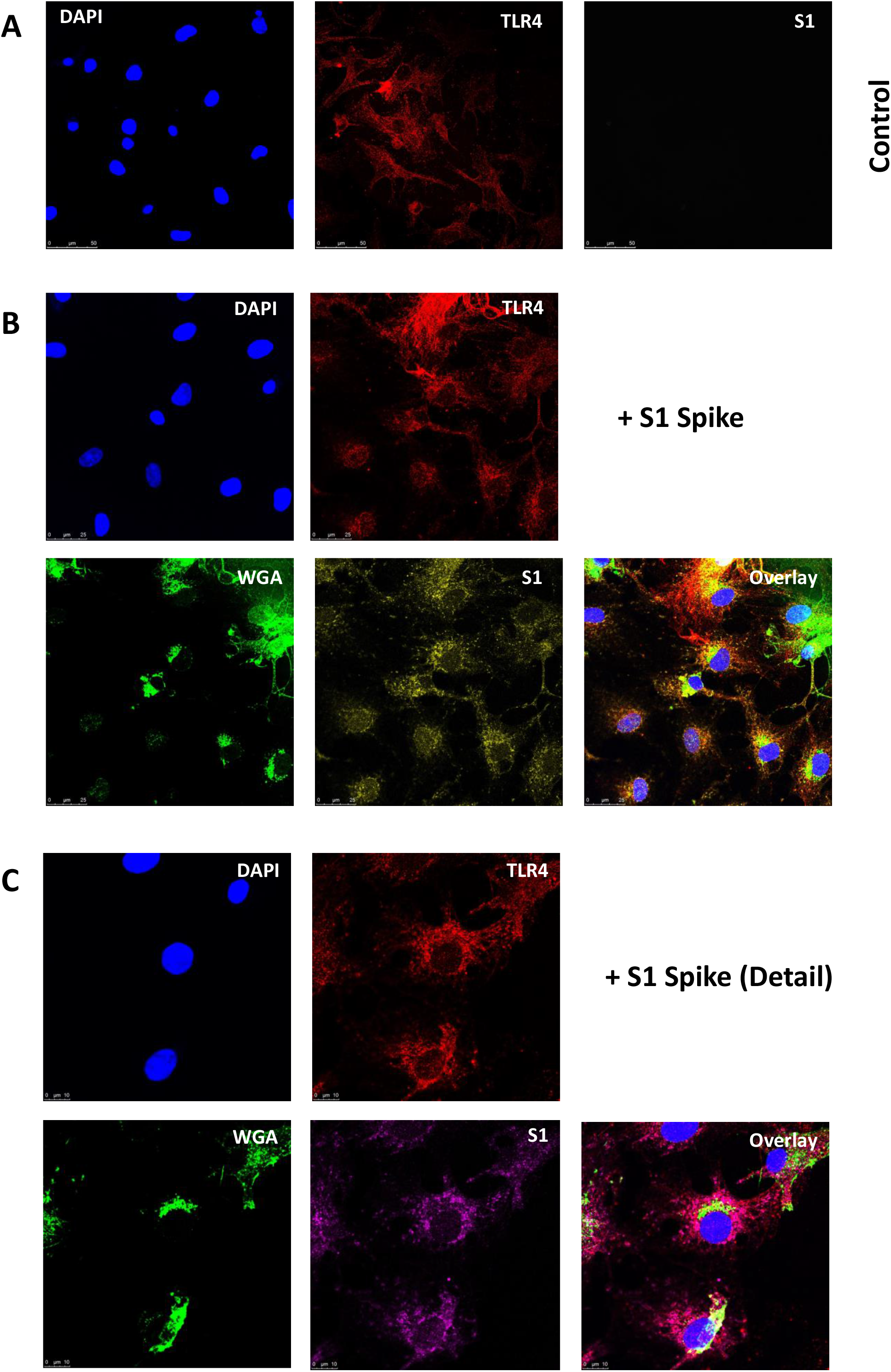
Confocal immunofluorescence analysis of S1 Spike-TLR4 co-localisation. **Panel A:** Control (untreated) cells (cTMFs) were grown on glass coverslips, fixed and stained using mouse monoclonal antibody against TLR4 or rabbit polyclonal antibody against S1 spike protein and corresponding fluorophore-conjugated secondary antibodies and counter-stained with the nuclear stain DAPI. Slides were then mounted and analysed using confocal immunofluorescence microscopy. TLR4/Cy™3 (red); S1/Alexa-Fluor® 647 (yellow); DAPI (blue). Magnification x63. Scale bar 25μm. **Panel B:** cTMFs were grown on glass coverslips and treated with recombinant S1 spike protein and 22 hours later fixed and stained using antibodies against TLR4 or S1 spike protein and corresponding fluorophore-conjugated secondary antibodies and counter-stained with the nuclear stain DAPI or the cellular membrane stain Alexa-Fluor® 488-conjugated wheat-germ agglutinin (WGA) [as for panel A: TLR4/Cy™3 (red); S1/Alexa-Fluor® 647 (yellow); DAPI (blue); WGA (green)]. Slides were then mounted and analysed using confocal immunofluorescence microscopy. Overlayed images are shown for comparison. Magnification x63. Scale bar 25μm. **Panel C:** As for panel B, but detail at x100 magnification. TLR4 (red); S1 (magenta); WGA (green); DAPI (blue).

**Figure 5.**
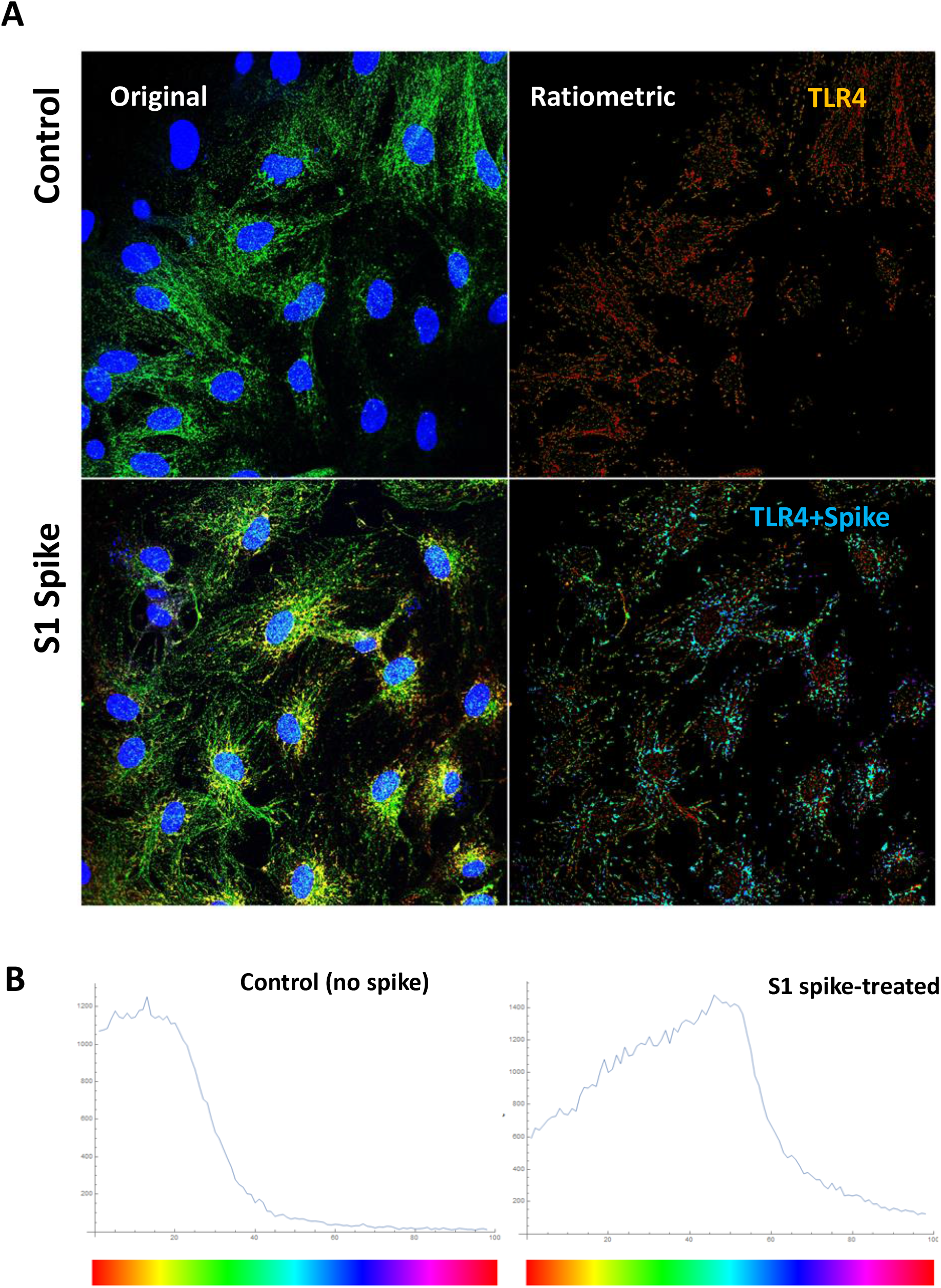
Ratiometric analysis of S1 spike and TLR4 co-localisation by confocal immunofluorescence. **Panel A:** Confocal images showing overlap of TLR4 and spike S1 in S1 treated cells or control images (no spike) were selected for ratiometric analysis of the degree of overlap between the fluorophores. TLR4 was imaged using Alexa-Fluor 488 and S1 spike using Alexa-Fluor® 647. Images were processed and a pseudo-colour scale applied in which red represents 0 or 1 at either end of the scale (no co-localisation) and cyan (centre) represents 0.5 (1:1 co-localisation). The weighted averages of the plotted data for control and spike were 0.245 and 0.438 respectively where 0.5 (cyan) represents an average 1:1 ratio of co-localisation. **Panel B:** Graphs showing the distribution of pixel intensity over the applied pseudo-colour spectrum for control (left) and spike S1-treated (right). On S1 spike treatment there was a marked shift towards the centre of the spectrum (cyan) approaching 1:1 co-localisation.

To confirm ACE2 upregulation as determined by immunoblotting, we treated cells with spike S1 protein and stained for ACE2. Figure 6 shows that the intensity of staining for ACE2 in the spike-treated cells, was higher than those in the control untreated cells, which agrees with the immunoblotting data shown in Figure 2.

**Figure 6.**
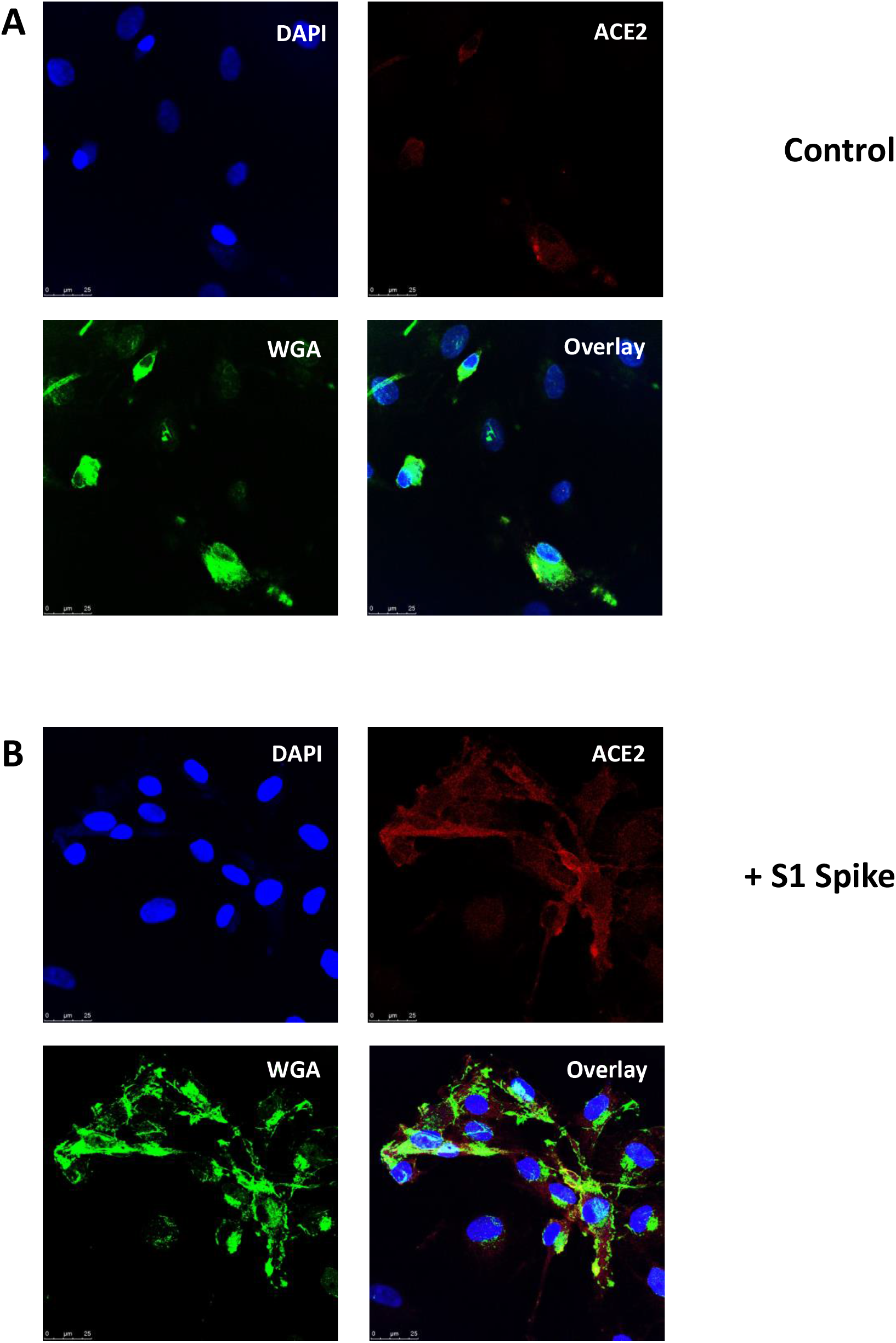
Confocal immunofluorescence analysis of ACE2 upregulation in S1 spike-treated cells. cTMFs were grown on glass coverslips and either left untreated or treated with recombinant S1 spike protein, fixed 22 hours later and stained using a rabbit polyclonal antibody against ACE2 and corresponding anti-rabbit Cy™3-conjugated secondary antibody (red). Samples were counter-stained with the nuclear stain DAPI (blue) or the cellular membrane stain Alexa-Fluor® 488-conjugated wheat-germ agglutinin (WGA: green). Slides were then mounted and analysed using confocal immunofluorescence microscopy. Magnification x63. Scale bar 25μm. **Panel A:** Control cells. **Panel B:** S1 spike-treated cells.

### Proximity ligation assay confirms binding of spike and TLR4 in rat and human cells

Cells were treated with spike S1 and then following fixation, permeabilization and blocking, each of TLR4 and spike S1 were probed with different primary antibodies. The Duolink® proximity ligation assay was used, which is a sensitive assay in which the respective secondary antibodies are conjugated to either plus or minus DNA strands, such that if the two proteins are in close proximity (at <40 nm resolution), a combination of DNA ligase and polymerase ligate and amplify the DNA to produce a fluorescence signal. Figure 7, panel B shows the red signal generated by amplified DNA corresponding to binding of both proteins in the spike S1-treated cTMFs but not in control cells (panel A). Following CLI-095 treatment, the red signals were more pronounced (results not shown), suggesting that preventing TLR4 internalisation increases binding on the cell surface. We then used human TLR4-HA HEK293 cells to confirm this result. Figure 8 shows that using either an anti-TLR4 antibody or an anti-HA tag antibody resulted in a strong PLA signal, confirming the TLR4-spike interaction.

**Figure 7.**
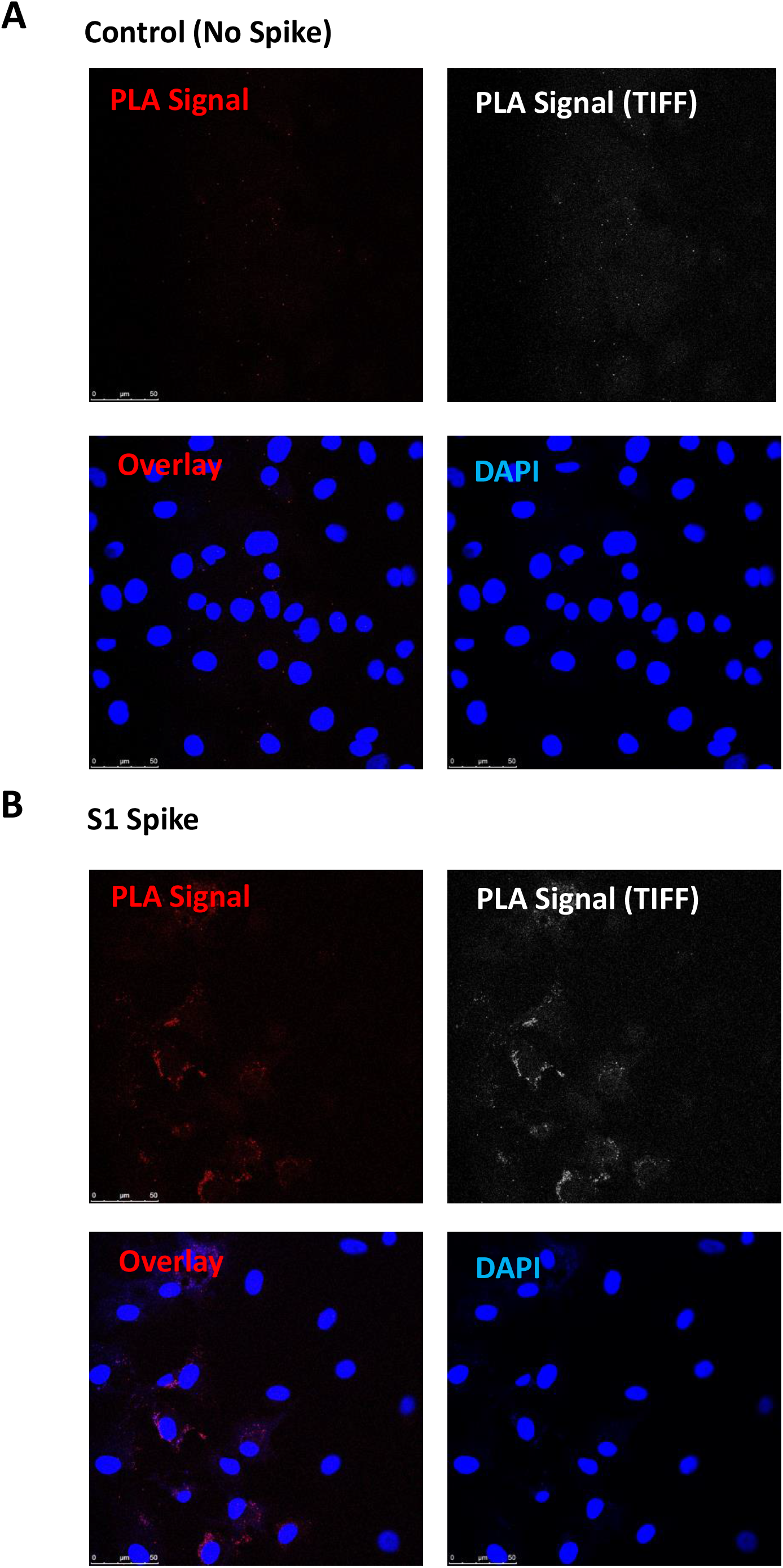
Proximity Ligation Assay of S1 spike-treated rat cardiac TM-derived fibrocytes (cTMFs). cTMFs were grown on coverslips and Proximity Ligation Assay (PLA) performed on control (no spike: **Panel A**) or S1 spike-treated cells (**Panel B**) with the Duolink® In Situ Red Mouse/Rabbit protocol using mouse anti-TLR4 and rabbit anti-spike S1 with the appropriate secondary antibodies (anti-rabbit-minus and anti-mouse-plus PLA probes). Coverslips were mounted in mounting medium containing DAPI and slides were analysed by confocal microscopy using the Alexa-Fluor® 594 and DAPI channels. A positive PLA signal appears as red dots or white in the corresponding TIFF image. Magnification x63. Scale bar 50μm.

**Figure 8.**
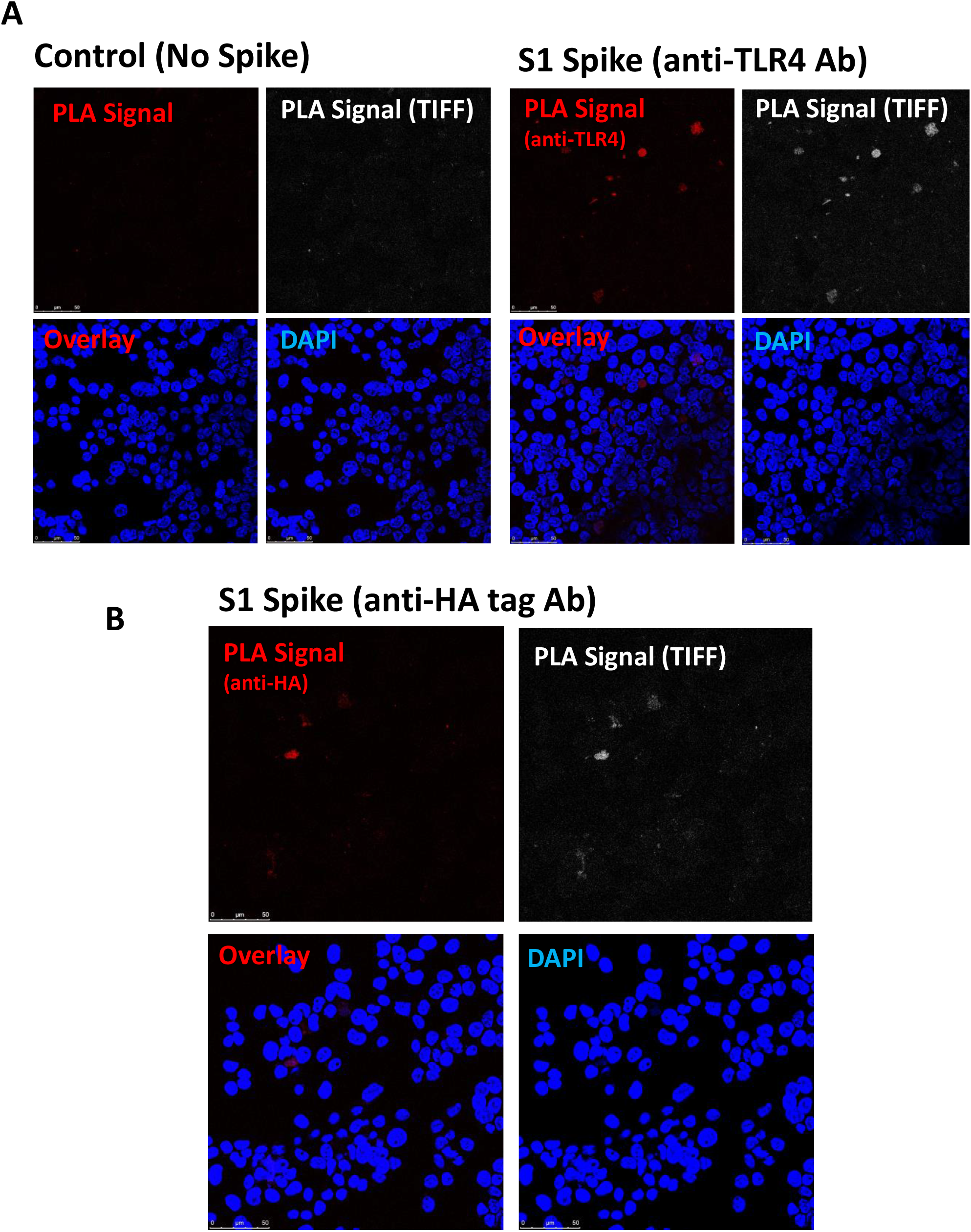
Proximity Ligation Assay of S1 spike-treated human TLR4-HA expressing HEK 293 cells. HEK 293 cells expressing HA-tagged human TLR4 were grown on coverslips and Proximity Ligation Assay (PLA) performed on control (no spike) or S1 spike-treated cells with the Duolink® In Situ Red Mouse/Rabbit protocol. Panel A: samples were probed using mouse anti-TLR4 and rabbit anti-spike S1 with the appropriate secondary antibodies (anti-rabbit-minus and anti-mouse-plus PLA probes). **Panel B:** samples were probed using mouse anti-HA and rabbit anti-spike S1. Coverslips were mounted in mounting medium containing DAPI and slides were analysed by confocal microscopy using the Alexa-Fluor® 594 and DAPI channels. A positive PLA signal appears as red dots or white in the corresponding TIFF image. Magnification x63. Scale bar 50μm.

### Spike S1 plus Interferon-γ induces a pro-inflammatory M1 macrophage phenotype in THP-1 monocyte-derived macrophages

THP-1 monocytes were differentiated into M_ø_ macrophages by PMA treatment for 48 hours. Following recovery in media, they were then treated with LPS + IFNγ as a positive control to induce M_1_ macrophage phenotype or spike S1+IFNγ, or IL-4 to induce M_2_ macrophage phenotype, or left untreated as controls. 24 hours after treatment, RNA was extracted for analysis by RT-qPCR. The results shown in Figure 9 demonstrate that spike S1+IFNγ also induced similar but weaker pro-inflammatory M_1_ macrophage polarisation in THP-1 cells, as illustrated by the marked increase in IL-1β and IL-6 transcript levels. A relatively low dose of spike S1, at 100ng/ml at which it optimally activates ACE2. Since spike S1 induces an M_1_ macrophage phenotype and inflammatory response at a dose lower than reported for ACE2 binding, this suggests that it could cause severe inflammation during viral sepsis, whether in the form of the intact virions or cleaved spike proteins circulating in the blood stream. CD206 (Mrc1) levels, as a marker for the pro-resolution M_2_ phenotype, were increased by IL-4 treatment but suppressed by M_1_ induction by both LPS+IFNγ and spike S1+IFNγ.

**Figure 9.**
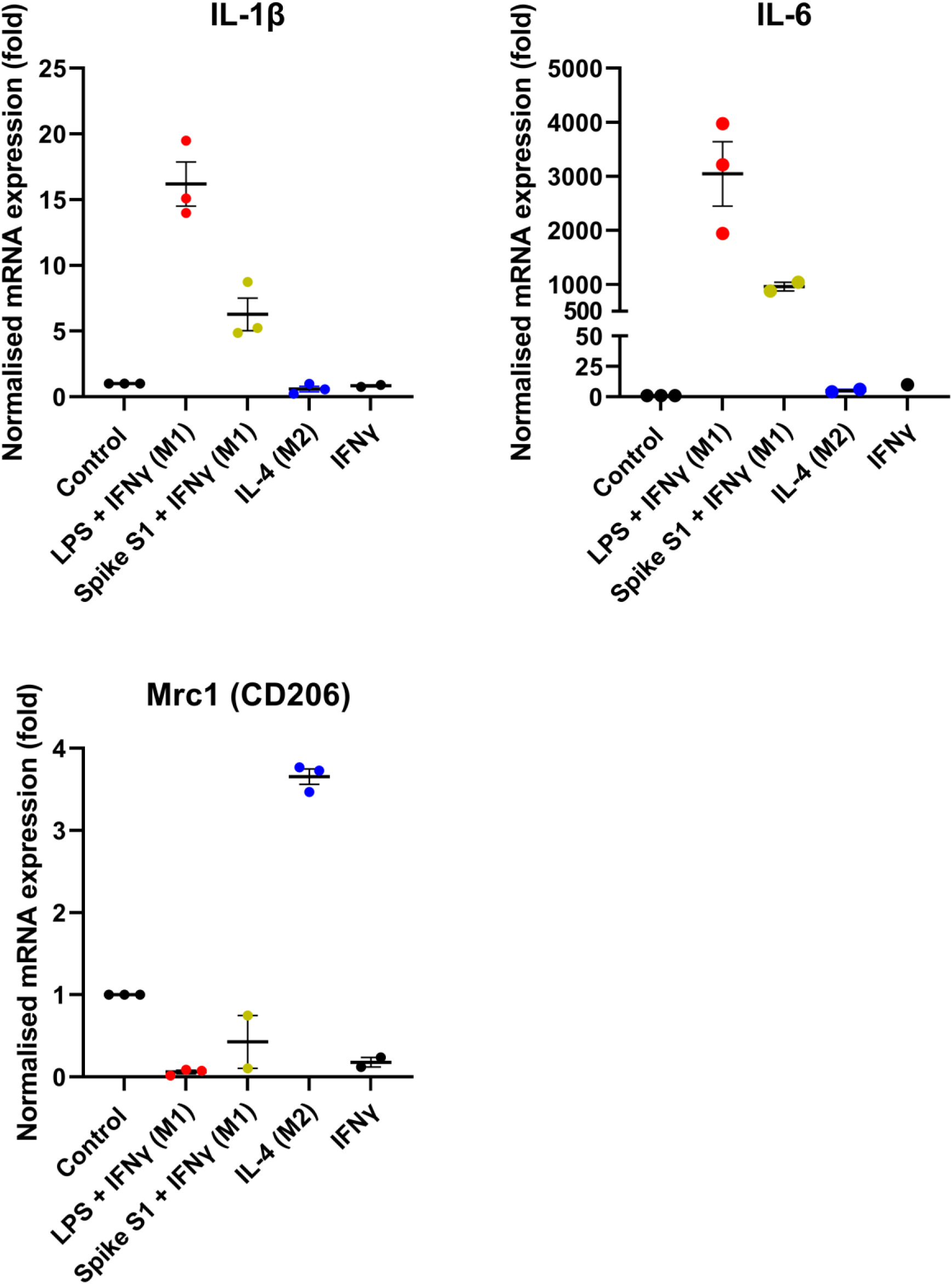
Analysis of M1 and M2 macrophage polarisation in LPS or S1 spike-treated human THP-1 monocyte-derived macrophages. Human THP-1 monocytic cells were differentiated into M_0_ macrophages using PMA and then further induced to M_1_ or M_2_ polarisation states suing LPS+interferon gamma (IFNγ) or interleukin-4 (IL-4) respectively, compared to cells treated with S1 spike+IFNγ or IFNγ alone. Following treatment RNA was isolated and analysed by RT-qPCR for IL-1β and IL-6 (M_1_) or Mrc1/Cd206 (M_2_).

## DISCUSSION

Here we show that the SARS-CoV-2 spike (S1) glycoprotein is a TLR4 agonist, which binds to and activates TLR4 to increase ACE2 expression within 22 hours in cardiac tissue-resident macrophage derived fibrocytes (cTMFs). The classical PAMP TLR4 agonist LPS also induced the same effect on ACE2 expression. Furthermore, the spike S1 downstream effects of induction of COX-2 expression, and downregulation of CTGF and Col3a1 are further evidence of a TLR4-mediated pathway, as indeed confirmed by inhibition by the TLR4 antagonist CLI-095. Increased ACE2 expression on the cell surface may facilitate SARS-CoV-2 virus entry and infectivity. As confirmed by Dynasore® inhibition, the TLR4 alternative TRIF-dependent endosomal pathway is responsible for the induction of ACE2 expression, which is consistent with the anti-inflammatory role of ACE2. As we have previously proposed, the intermediate mechanism is likely mediated via type I interferon production acting in an autocrine or paracrine manner [1], since ACE2 is an interferon-stimulated gene (ISG) [54], although the exact intermediate link remains to be confirmed in these cells. Interestingly, since we originally proposed this mechanism, SARS-CoV-2 spike protein has been shown to bind to LPS and that this interaction augments NFκB activation and cytokine expression in human monocytic THP-1 cells and peripheral blood mononuclear cells (PBMCs) *in vitro* and also *in vivo* in NFκB reporter mice [55].

Together these findings suggest a mechanism for the increased virulence of SARS-CoV-2, since TLR4 activation upregulates ACE2 expression in cells which otherwise have relatively low levels of ACE2, demonstrating that this is a generic effect of TLR4 activation by PAMPs and VAMPs. Furthermore, TLR4 activation leads to pro-inflammatory cytokine release. There are likely two scenarios which contribute to these effects: (i) the SARS-CoV-2 virions themselves bind to TLR4 via their spike glycoproteins to increase ACE2 expression and hence promote infectivity and (ii) the cleaved S1 glycoprotein subunits which are released into the interstitial space and blood stream would activate TLR4 in tissues. The outer S1 subunits are cleaved from spike by furin/TMPRSS2 or cathepsins B/L in the endosomes [37], following cell engagement with ACE2 for example, during the viral entry process [27, 34]. However, whilst this generalised model is true for SARS-CoV and SARS-CoV-2 [33-36], in fact for SARS-CoV-2 the story may be more complicated because furin-dependent cleavage of newly synthesised spike protein also occurs in the trans-Golgi complex due to a unique 4 amino-acid insert within the spike protein at the S1-S2 junction [24, 31]. This means that cleaved spike S1 may also activate TLR4 intracellularly (e.g. possibly in endosomes), but more importantly it would be exported out of the cell in the secretory pathway during the viral exit process [32, 56]. Furthermore, there is also evidence for shedding of the S1 domain into the extracellular space where it could activate TLR4, especially since spike S1 was detected in both the blood and urine of COVID-19 patients [57, 58]. Indeed, the presence of this furin cleavage site has been shown to promote viral entry into lung cells and contribute to pathogenesis, although it is not essential for infection [28, 33, 37, 59]. Based on our novel findings, we hereby unravel part of the mechanisms responsible for this; the higher numbers of cleaved S1 at the cell surface (or in endosomes) during viral entry or exported out of the infected cell would activate TLR4 on neighbouring cells to increase ACE2 expression thereby promoting cellular entry and also causing hyperinflammation, both of which may be key to the increased infectivity, virulence, tissue tropism and mortality associated with SARS-CoV-2. Importantly, there is also some evidence that the cleaved S1 and S2 subunits can remain bound non-covalently, in a “metastable pre-fusion state” which possibly facilitates infectivity/tropism in host cells which lack TMPRSS2 [24, 32, 37, 60].

### TLR4 as a possible entry receptor/co-factor

ACE2 expression is mediated via the TLR4 alternative endosomal pathway, since it was inhibited by Dynasore®, a dynamin inhibitor. This suggests that the spike S1-TLR4 complex is internalised in the cell into endosomes, and therefore TLR4 may facilitate entry of SARS-CoV-2 by acting as a modulator/co-factor for entry via the endosomal pathway. Receptor-mediated, clathrin-dependent endocytosis is a key mechanism for viral entry into host cells, including SARS-CoV-1 [61] and SARS-CoV-2 (via ACE2 or otherwise) [27, 62, 63]. The endocytic pathway has also been the target for some proposed drugs for COVID-19 [27, 62, 63]. The same endocytic mechanisms apply for TLR4 endocytosis; and since spike S1 binds TLR4 and is internalised into endosomes via receptor-mediated endocytosis based on our findings, then TLR4 could represent an additional route of viral entry, either independently of ACE2 or via ACE2.

In fact, TLR4 is involved in the internalisation of Gram-negative E. coli bacteria into phagosomes, via its adaptor TRAM interacting with the Rab11-family interacting protein 2 (Rab11-FIP2). [64, 65]. In terms of signalling, Rab11a GTPase also controls TLR4-induced stimulation of IRF3 on phagosomes [65].

Immunocytochemistry and proximity ligation assays strongly suggest that spike is binding TLR4. This also indicates that spike is still present in these cells after almost 22 hours and the TLR4-spike S1 complex is therefore probably being internalised into the cell in endosomes. Subsequent cleavage of spike via endosomal cathepsins B and L may therefore facilitate viral entry, meaning that TLR4-mediated internalisation into endosomes positively facilitates and/or modulates SARS-CoV-2 viral entry [37].

### Effects of Dynasore® on COX-2 expression

Whereas COX-2 induction by LPS and spike S1 was completely abrogated by CLI-095, it was essentially unaffected by dynamin inhibition with Dynasore® in the case of LPS. This suggests that COX-2 expression is not dependent on the alternative TRIF-dependent endosomal pathway. This suggests that either COX-2 induction is dependent on the canonical MyD88 pathway since the COX-2 proximal gene promoter region has both NFκB and NFAT transcription sites [9] and some studies have shown it to be regulated by the canonical MyD88-dependent pathway in peritoneal macrophages [48]. However, it is also consistent with our previous findings that in cTMFs, COX-2 induction downstream of TLR4 activation is mediated via the CN/NFAT and PKCε pathway and not NFκB [9].

The observation of an enhanced COX-2 expression with spike S1 plus Dynasore®, compared to spike S1 alone, supports a related phenomenon: after the initial TLR4 stimulation, receptor-mediated endocytosis itself serves as a negative regulator of LPS-TLR4 signalling. Blocking this via Dynasore®, keeps TLR4 at the cell surface while also inhibiting the downstream negative feedback loops of the alternative pathway and expression of feedback inhibitors, so there is an increase in canonical, unopposed, MyD88-NFκB or Ca^2+^/CN/NFAT proinflammatory signalling. This idea is illustrated by the fact that inhibition of endocytosis of the IL-1 receptor (which shares the TIR domain with TLRs) by Dynasore®, disrupts the feedback inhibition loops on mitogen-activated protein kinase and NFκB signalling (although the pathway loop is more complex for the latter) [66]. COX-2 (Ptgs2) mRNA expression increased approx. 3.7 fold in the presence of Dynasore® after IL-1 stimulation at 60 min, and was NFκB-dependent [66].

Thus, inhibition of the endosomal pathway inhibits the negative feedback on the canonical NFκB-dependent expression of genes. Furthermore, many other molecules exert negative feedback regulation on TLR4 signalling, (See [67] and [18] for reviews). Moreover, TRAF3 ubiquitination states in the alternative pathway selectively regulate the expression of type I interferons and proinflammatory cytokines [68]. Thus, it appears that the spike S1 balance of biased TLR4 signalling towards one pathway or another, downstream of TLR4, may be slightly different from LPS. Collectively, these findings implicate TLR4 activation as one of the possible drivers of the COVID-19 hyperinflammation and “cytokine storm”.

### Dynasore® effects on CTGF and Col3a1 expression

Dynasore® appeared to enhance the downregulation of CTGF expression by LPS and spike S1. Dynamin is not only important for endocytosis but also in fusion pore expansion during exocytosis (as the vesicle fuses with the plasma membrane), as well as in intracellular transport, budding from Golgi apparatus and ER and cell division (see [52, 53, 69-71]). Since CTGF and Col3a1 are secreted proteins, their intracellular transport via budding vesicles from the ER and Golgi apparatus, as well as their secretion would be inhibited by Dynasore® which could contribute to the enhanced downregulation. This is in line with evidence that Dynasore inhibits MMP-3 induced CTGF/CCN2 protein expression in human dental pulp cells (fibroblast-like cells) [72]. However, we cannot exclude gene regulatory effects on CTGF/Collagen expression at the transcriptional level, as suggested by our previous study [9] and the fact that total levels, as detected by immunoblotting of whole cell extracts, reflect overall expression levels rather than subtle differences in trafficking through different intracellular compartments.

### Translational therapeutic implications for treatment of COVID-19: TLR4 antagonists

Our findings represent a potentially important advance in deciphering the pathophysiological mechanisms occurring in COVID-19 as well as providing evidence for the use of TLR4 antagonists for its treatment (see also [1]). TLR4 antagonists, particularly Resatorvid®, will be useful therapeutically in the later severe stages particularly to reduce or prevent the excessive TLR4-mediated hyperinflammation, or cytokine storm. We propose the clinical trial testing of Resatorvid® (also known as CLI-095 or TAK-242) as a treatment in patients with COVID-19 disease associated with hyperinflammation and/or multi-organ involvement, to reduce the susceptibility of cells to further infection and reduce the inflammatory symptoms.

Our results also suggest that macrophages, particularly tissue resident macrophages, and dendritic cells may be a target cell for SARS-CoV-2, especially given that they express high levels of TLR4. Furthermore, the balance of activation of the canonical vs. alternative pathway may follow spatial (i.e. which tissue) and temporal (i.e. time of activation depending on course of infection) variation. Therefore, the activation of TLR4 by the spike S1 domain of SARS-CoV-2 virus may cause hyperinflammation via the canonical pathway and/or increase ACE2 expression via the alternative endosomal pathway, thus increasing viral entry via ACE2. A schematic representation of the mechanisms resulting in ACE2 upregulation, increased viral entry and inflammation are shown in Figure 10.

**Figure 10.**
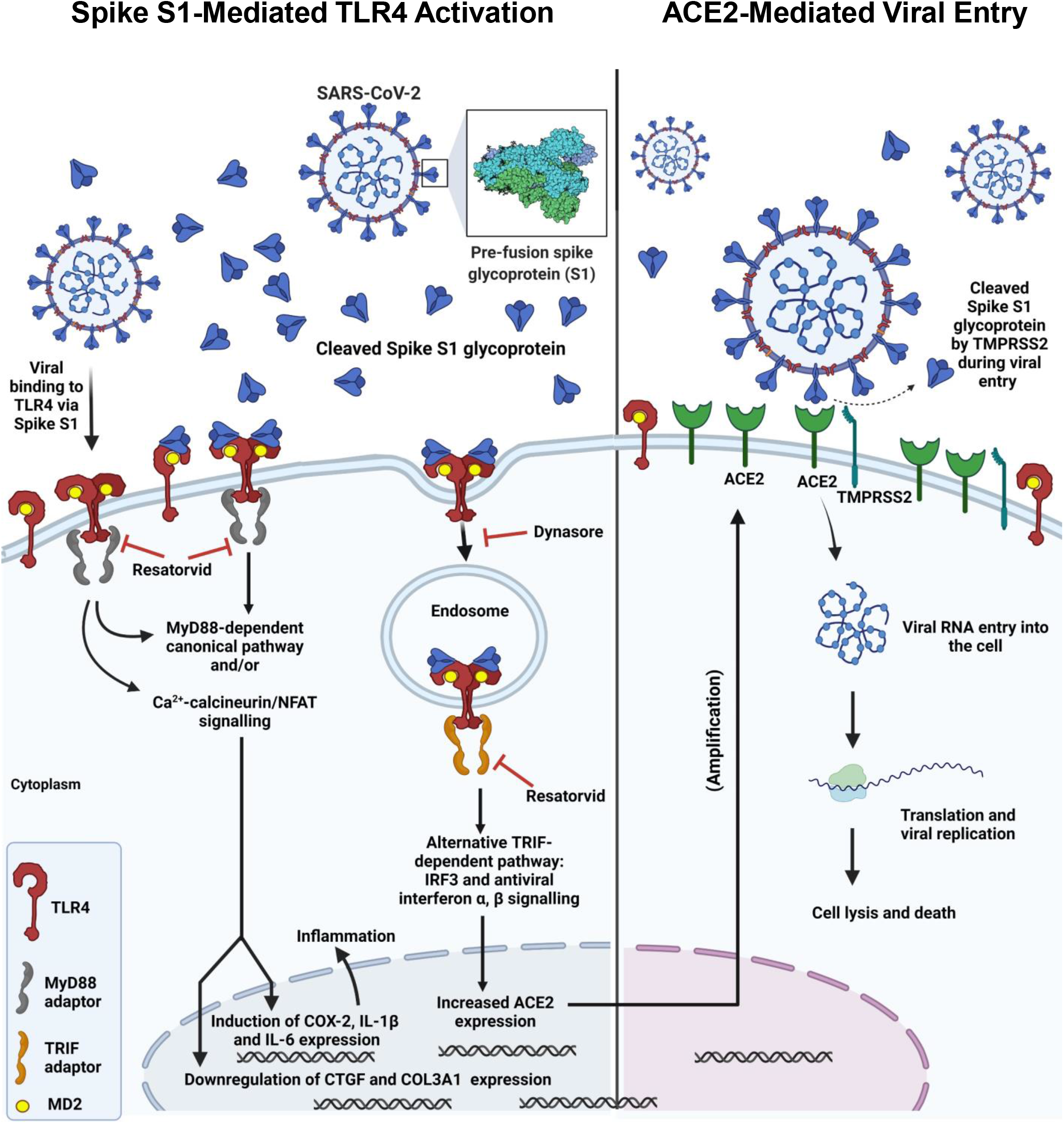
S1 spike-mediated TLR4 activation leading to inflammation, ACE2 upregulation and increased viral entry. Schematic showing SARS-CoV-2 spike interaction with TLR4, subsequent TLR4 activation via canonical and endosomal pathways and induction of inflammation, upregulation of ACE2 expression and potential increased viral entry. Created with BioRender.com™.

Patients with concomitant gram-negative bacterial infections are more susceptible to SARS-CoV-2 infections, because LPS increases ACE2 expression. LPS also increases TLR4 expression but through the canonical NFκB pathway (see [1] and references within, and [73]) Furthermore, spike S1 binds to LPS increasing TLR-mediated proinflammatory cytokine expression [55]. Patients with cardiovascular comorbidities could be more susceptible to SARS-CoV-2 infections not only because they have more ACE2 expression, but also because they have increased cardiovascular TLR4 expression and activation and therefore basal inflammation.

## CONCLUSIONS

We have shown that SARS-CoV-2 spike S1 glycoprotein is a TLR4 agonist. We have also shown that spike S1, like LPS, activates TLR4 and increases ACE2 expression in adult rat cardiac tissue resident macrophage-derived fibrocytes/fibroblasts and human THP-1 monocyte-derived macrophages. This novel finding is important because the mechanism may facilitate viral entry into cardiac and other cells which otherwise have low ACE2 expression. This effect of ACE2 upregulation is blocked by the specific TLR4 signalling inhibitor CLI-095 (Resatorvid®), which reduced ACE2 expression to basal levels. ACE2 upregulation is induced via the dynamin-dependent endosomal pathway, indicating that spike S1-TLR4 complex is internalised into endosomes to activate the alternative TRIF/TRAM pathway signalling. Spike S1 also led to COX-2 induction and marked downregulation of CTGF and COL3A1 very similar to the effects conferred by Gram negative bacterial LPS. The spike S1 effects on COX-2 induction were also inhibited by CLI-095 (Resatorvid®). Spike S1 in conjunction with IFN-γ was able to induce pro-inflammatory M_1_ macrophage polariation and expression of the pro-inflammatory cytokines IL-1 β and IL-6. Collectively, these findings also suggest that TLR4 may be a possible entry receptor or co-factor for the virus. We suggest the need for clinical trial testing of Resatorvid® and perhaps other TLR4 antagonists for the treatment of severe COVID-19 disease, especially given that it was well tolerated in a previous clinical trial for sepsis, apart from a minor reversible side effect of methemoglobinemia [74]. Resatorvid® will prevent ACE2 upregulation, reduce cellular susceptibility to viral infection and reduce the hyperinflammation occurring in COVID-19. In effect, it will serve as an antiviral and anti-inflammatory drug.

## Acknowledgements

We thank Dr Rhys Anderson (Cardiovascular Research Section, King’s College London) for help with the Duolink® Proximity Ligation Assay and Vasco Claro (Vascular Biology and Inflammation, King’s College London) for help setting up RT-qPCR and primers for M_1_ and M_2_ macrophage markers.

M.M.A. is funded by a King’s International Postgraduate Research Scholarship.

## Conflict of interests

The authors declare that they have no conflict of interest.

## Notes

### Competing Interest Statement

The authors have declared no competing interest.

## REFERENCES

1. Aboudounya, M.M. and R.J. Heads, COVID-19 and Toll-Like Receptor 4 (TLR4): SARS-CoV-2 May Bind and Activate TLR4 to Increase ACE2 Expression, Facilitating Entry and Causing Hyperinflammation. Mediators Inflamm, 2021. 2021: p. 8874339.

2. Hoffmann, M., et al., SARS-CoV-2 Cell Entry Depends on ACE2 and TMPRSS2 and Is Blocked by a Clinically Proven Protease Inhibitor. Cell, 2020. 181(2): p. 271–280 e8.

3. Wan, Y., et al., Receptor Recognition by the Novel Coronavirus from Wuhan: an Analysis Based on Decade-Long Structural Studies of SARS Coronavirus. J Virol, 2020. 94(7).

4. Zou, X., et al., Single-cell RNA-seq data analysis on the receptor ACE2 expression reveals the potential risk of different human organs vulnerable to 2019-nCoV infection. Front Med, 2020. 14(2): p. 185–192.

5. Fu, J., et al., Expressions and significances of the angiotensin-converting enzyme 2 gene, the receptor of SARS-CoV-2 for COVID-19. Mol Biol Rep, 2020. 47(6): p. 4383–4392.

6. Hikmet, F., et al., The protein expression profile of ACE2 in human tissues. Mol Syst Biol, 2020. 16(7): p. e9610.

7. Tufan, A., A. Avanoglu Guler, and M. Matucci-Cerinic, COVID-19, immune system response, hyperinflammation and repurposing antirheumatic drugs. Turk J Med Sci, 2020. 50(SI-1): p. 620–632.

8. Akira, S., S. Uematsu, and O. Takeuchi, Pathogen recognition and innate immunity. Cell, 2006. 124(4): p. 783–801.

9. Mesquita, R.F., et al., Protein kinase Cepsilon-calcineurin cosignaling downstream of toll-like receptor 4 downregulates fibrosis and induces wound healing gene expression in cardiac myofibroblasts. Mol Cell Biol, 2014. 34(4): p. 574–94.

10. Armstrong, L., et al., Expression of functional toll-like receptor-2 and -4 on alveolar epithelial cells. Am J Respir Cell Mol Biol, 2004. 31(2): p. 241–5.

11. Molteni, M., S. Gemma, and C. Rossetti, The Role of Toll-Like Receptor 4 in Infectious and Noninfectious Inflammation. Mediators Inflamm, 2016. 2016: p. 6978936.

12. Satoh, T. and S. Akira, Toll-Like Receptor Signaling and Its Inducible Proteins. Microbiol Spectr, 2016. 4(6).

13. Husebye, H., et al., Endocytic pathways regulate Toll-like receptor 4 signaling and link innate and adaptive immunity. EMBO J, 2006. 25(4): p. 683–92.

14. Schoggins, J.W. and C.M. Rice, Interferon-stimulated genes and their antiviral effector functions. Curr Opin Virol, 2011. 1(6): p. 519–25.

15. Mogensen, T.H. and S.R. Paludan, Reading the viral signature by Toll-like receptors and other pattern recognition receptors. J Mol Med (Berl), 2005. 83(3): p. 180–92.

16. Barton, G.M. and R. Medzhitov, Toll-like receptor signaling pathways. Science, 2003. 300(5625): p. 1524–5.

17. Bhattacharyya, S., et al., Toll-Like Receptor-4 Signaling Drives Persistent Fibroblast Activation and Prevents Fibrosis Resolution in Scleroderma. Adv Wound Care (New Rochelle), 2017. 6(10): p. 356–369.

18. Yang, Y., et al., The emerging role of Toll-like receptor 4 in myocardial inflammation. Cell Death Dis, 2016. 7: p. e2234.

19. Akira, S. and K. Takeda, Toll-like receptor signalling. Nat Rev Immunol, 2004. 4(7): p. 499–511.

20. Chow, J.C., et al., Toll-like receptor-4 mediates lipopolysaccharide-induced signal transduction. J Biol Chem, 1999. 274(16): p. 10689–92.

21. Akashi, S., et al., Lipopolysaccharide interaction with cell surface Toll-like receptor 4-MD-2: higher affinity than that with MD-2 or CD14. J Exp Med, 2003. 198(7): p. 1035–42.

22. Marongiu, L., et al., Below the surface: The inner lives of TLR4 and TLR9. J Leukoc Biol, 2019. 106(1): p. 147–160.

23. Wang, Y., et al., Inhibition of clathrin/dynamin-dependent internalization interferes with LPS-mediated TRAM-TRIF-dependent signaling pathway. Cell Immunol, 2012. 274(1-2): p. 121–9.

24. Walls, A.C., et al., Structure, Function, and Antigenicity of the SARS-CoV-2 Spike Glycoprotein. Cell, 2020. 183(6): p. 1735.

25. Wrapp, D., et al., Cryo-EM structure of the 2019-nCoV spike in the prefusion conformation. Science, 2020. 367(6483): p. 1260–1263.

26. Choudhury, A. and S. Mukherjee, In silico studies on the comparative characterization of the interactions of SARS-CoV-2 spike glycoprotein with ACE-2 receptor homologs and human TLRs. J Med Virol, 2020.

27. Al-Horani, R.A., S. Kar, and K.F. Aliter, Potential Anti-COVID-19 Therapeutics that Block the Early Stage of the Viral Life Cycle: Structures, Mechanisms, and Clinical Trials. Int J Mol Sci, 2020. 21(15).

28. Hoffmann, M., H. Kleine-Weber, and S. Pohlmann, A Multibasic Cleavage Site in the Spike Protein of SARS-CoV-2 Is Essential for Infection of Human Lung Cells. Mol Cell, 2020. 78(4): p. 779–784 e5.

29. Yuan, M., et al., A highly conserved cryptic epitope in the receptor binding domains of SARS-CoV-2 and SARS-CoV. Science, 2020. 368(6491): p. 630–633.

30. Bestle, D., et al., TMPRSS2 and furin are both essential for proteolytic activation of SARS-CoV-2 in human airway cells. Life Sci Alliance, 2020. 3(9).

31. Jaimes, J.A., J.K. Millet, and G.R. Whittaker, Proteolytic Cleavage of the SARS-CoV-2 Spike Protein and the Role of the Novel S1/S2 Site. iScience, 2020. 23(6): p. 101212.

32. Duan, L., et al., The SARS-CoV-2 Spike Glycoprotein Biosynthesis, Structure, Function, and Antigenicity: Implications for the Design of Spike-Based Vaccine Immunogens. Front Immunol, 2020. 11: p. 576622.

33. Papa, G., et al., Furin cleavage of SARS-CoV-2 Spike promotes but is not essential for infection and cell-cell fusion. PLoS Pathog, 2021. 17(1): p. e1009246.

34. Xia, S., et al., The role of furin cleavage site in SARS-CoV-2 spike protein-mediated membrane fusion in the presence or absence of trypsin. Signal Transduct Target Ther, 2020. 5(1): p. 92.

35. Glowacka, I., et al., Evidence that TMPRSS2 activates the severe acute respiratory syndrome coronavirus spike protein for membrane fusion and reduces viral control by the humoral immune response. J Virol, 2011. 85(9): p. 4122–34.

36. Belouzard, S., V.C. Chu, and G.R. Whittaker, Activation of the SARS coronavirus spike protein via sequential proteolytic cleavage at two distinct sites. Proc Natl Acad Sci U S A, 2009. 106(14): p. 5871–6.

37. Evans, J.P. and S.L. Liu, Role of host factors in SARS-CoV-2 entry. J Biol Chem, 2021. 297(1): p. 100847.

38. Braun, E. and D. Sauter, Furin-mediated protein processing in infectious diseases and cancer. Clin Transl Immunology, 2019. 8(8): p. e1073.

39. Coutard, B., et al., The spike glycoprotein of the new coronavirus 2019-nCoV contains a furin-like cleavage site absent in CoV of the same clade. Antiviral Res, 2020. 176: p. 104742.

40. Wu, C., et al., Furin: A Potential Therapeutic Target for COVID-19. iScience, 2020. 23(10): p. 101642.

41. Brunton, B., et al., TIM-1 serves as a receptor for Ebola virus in vivo, enhancing viremia and pathogenesis. PLoS Negl Trop Dis, 2019. 13(6): p. e0006983.

42. Dragovich, M.A., et al., Biomechanical characterization of TIM protein-mediated Ebola virus-host cell adhesion. Sci Rep, 2019. 9(1): p. 267.

43. Chanput, W., J.J. Mes, and H.J. Wichers, THP-1 cell line: an in vitro cell model for immune modulation approach. Int Immunopharmacol, 2014. 23(1): p. 37–45.

44. Matsunaga, N., et al., TAK-242 (resatorvid), a small-molecule inhibitor of Toll-like receptor (TLR) 4 signaling, binds selectively to TLR4 and interferes with interactions between TLR4 and its adaptor molecules. Mol Pharmacol, 2011. 79(1): p. 34–41.

45. Takashima, K., et al., Analysis of binding site for the novel small-molecule TLR4 signal transduction inhibitor TAK-242 and its therapeutic effect on mouse sepsis model. Br J Pharmacol, 2009. 157(7): p. 1250–62.

46. Kagan, J.C., et al., TRAM couples endocytosis of Toll-like receptor 4 to the induction of interferon-beta. Nat Immunol, 2008. 9(4): p. 361–8.

47. Macia, E., et al., Dynasore, a cell-permeable inhibitor of dynamin. Dev Cell, 2006. 10(6): p. 839–50.

48. Kawai, T., et al., Lipopolysaccharide stimulates the MyD88-independent pathway and results in activation of IFN-regulatory factor 3 and the expression of a subset of lipopolysaccharide-inducible genes. J Immunol, 2001. 167(10): p. 5887–94.

49. Hirotani, T., et al., Regulation of lipopolysaccharide-inducible genes by MyD88 and Toll/IL-1 domain containing adaptor inducing IFN-beta. Biochem Biophys Res Commun, 2005. 328(2): p. 383–92.

50. Zanoni, I., et al., CD14 regulates the dendritic cell life cycle after LPS exposure through NFAT activation. Nature, 2009. 460(7252): p. 264–8.

51. Ryu, J.K., et al., Reconstruction of LPS Transfer Cascade Reveals Structural Determinants within LBP, CD14, and TLR4-MD2 for Efficient LPS Recognition and Transfer. Immunity, 2017. 46(1): p. 38–50.

52. Anantharam, A., et al., A new role for the dynamin GTPase in the regulation of fusion pore expansion. Mol Biol Cell, 2011. 22(11): p. 1907–18.

53. Preta, G., J.G. Cronin, and I.M. Sheldon, Dynasore - not just a dynamin inhibitor. Cell Commun Signal, 2015. 13: p. 24.

54. Ziegler, C.G.K., et al., SARS-CoV-2 Receptor ACE2 Is an Interferon-Stimulated Gene in Human Airway Epithelial Cells and Is Detected in Specific Cell Subsets across Tissues. Cell, 2020. 181(5): p. 1016–1035 e19.

55. Petruk, G., et al., SARS-CoV-2 spike protein binds to bacterial lipopolysaccharide and boosts proinflammatory activity. J Mol Cell Biol, 2020. 12(12): p. 916–932.

56. Belouzard, S., et al., Mechanisms of coronavirus cell entry mediated by the viral spike protein. Viruses, 2012. 4(6): p. 1011–33.

57. George, S., et al., Evidence for SARS-CoV-2 Spike Protein in the Urine of COVID-19 Patients. Kidney360, 2021. 2(6): p. 922–934.

58. Ogata, A.F., et al., Ultra-Sensitive Serial Profiling of SARS-CoV-2 Antigens and Antibodies in Plasma to Understand Disease Progression in COVID-19 Patients with Severe Disease. Clin Chem, 2020. 66(12): p. 1562–1572.

59. Johnson, B.A., et al., Loss of furin cleavage site attenuates SARS-CoV-2 pathogenesis. Nature, 2021. 591(7849): p. 293–299.

60. Kirchdoerfer, R.N., et al., Pre-fusion structure of a human coronavirus spike protein. Nature, 2016. 531(7592): p. 118–21.

61. Wang, H., et al., SARS coronavirus entry into host cells through a novel clathrin-and caveolae-independent endocytic pathway. Cell Res, 2008. 18(2): p. 290–301.

62. Glebov, O.O., Understanding SARS-CoV-2 endocytosis for COVID-19 drug repurposing. FEBS J, 2020. 287(17): p. 3664–3671.

63. Yang, N. and H.M. Shen, Targeting the Endocytic Pathway and Autophagy Process as a Novel Therapeutic Strategy in COVID-19. Int J Biol Sci, 2020. 16(10): p. 1724–1731.

64. Skjesol, A., et al., The TLR4 adaptor TRAM controls the phagocytosis of Gram-negative bacteria by interacting with the Rab11-family interacting protein 2. PLoS Pathog, 2019. 15(3): p. e1007684.

65. Husebye, H., et al., The Rab11a GTPase controls Toll-like receptor 4-induced activation of interferon regulatory factor-3 on phagosomes. Immunity, 2010. 33(4): p. 583–96.

66. Hansen, B., et al., Regulation of NF-kappaB-dependent gene expression by ligand-induced endocytosis of the interleukin-1 receptor. Cell Signal, 2013. 25(1): p. 214–28.

67. Anwar, M.A., S. Basith, and S. Choi, Negative regulatory approaches to the attenuation of Toll-like receptor signaling. Exp Mol Med, 2013. 45: p. e11.

68. Tseng, P.H., et al., Different modes of ubiquitination of the adaptor TRAF3 selectively activate the expression of type I interferons and proinflammatory cytokines. Nat Immunol, 2010. 11(1): p. 70–5.

69. Gonzalez-Jamett, A.M., et al., Dynamin-2 regulates fusion pore expansion and quantal release through a mechanism that involves actin dynamics in neuroendocrine chromaffin cells. PLoS One, 2013. 8(8): p. e70638.

70. Shin, W., et al., Visualization of Membrane Pore in Live Cells Reveals a Dynamic-Pore Theory Governing Fusion and Endocytosis. Cell, 2018. 173(4): p. 934–945 e12.

71. Jackson, J., et al., Small molecules demonstrate the role of dynamin as a bi-directional regulator of the exocytosis fusion pore and vesicle release. Mol Psychiatry, 2015. 20(7): p. 810–9.

72. Muromachi, K., et al., MMP-3 provokes CTGF/CCN2 production independently of protease activity and dependently on dynamin-related endocytosis, which contributes to human dental pulp cell migration. J Cell Biochem, 2012. 113(4): p. 1348–58.

73. Wang, P., et al., LPS enhances TLR4 expression and IFNgamma production via the TLR4/IRAK/NFkappaB signaling pathway in rat pulmonary arterial smooth muscle cells. Mol Med Rep, 2017. 16(3): p. 3111–3116.

74. Rice, T.W., et al., A randomized, double-blind, placebo-controlled trial of TAK-242 for the treatment of severe sepsis. Crit Care Med, 2010. 38(8): p. 1685–94.

